# Beyond two alleles: Multiallelic genotypic selection and its estimation from time-series data

**DOI:** 10.1101/2024.11.08.622587

**Authors:** Nikolas Vellnow, Toni I. Gossmann, David Waxman

## Abstract

Genetic diversity is central to evolutionary change, with both natural selection and random genetic drift depending on variation within a population. An individual in a diploid population carries two alleles per locus, yet the population as a whole can harbour many alleles, giving rise to a rich spectrum of homozygous and heterozygous genotypes. Such multiallelic variation is common at biologically and medically important loci such as the major histocompatibility complex, the ABO blood group system, and genes underlying monogenic diseases. However, much of population genetic theory and data analysis has focussed on biallelic loci.

Here, we introduce a matrix representation of the genotypic selection acting at a multiallelic locus. This exploits the common mathematical structure underlying selection and drift, and separates the effects of genetic diversity and fitness. The representation accommodates diverse selection regimes, including additive, multiplicative, frequency-dependent, and temporally varying selection, as well as heterozygote advantage. We show how, under specific assumptions, genotype-specific fitness-effects can be estimated from allele frequency trajectories over microevolutionary timescales. Applying this estimation procedure to time-series data from experimental yeast evolution illustrates how multiallelic fitness interactions, including heterozygote advantage, may be characterised from haplotype frequency data. More broadly, this work provides a practical foundation for analysing evolutionary dynamics at multiallelic loci in experimental and natural populations.

## Introduction

Genetic diversity plays a central role in evolutionary theory, as it provides the raw material on which natural selection and random genetic drift act. In population genetics, diversity is most commonly quantified at the level of single nucleotides or loci, with a predominant focus on *biallelic* single nucleotide polymorphisms (SNPs) (Heyne et al., 2023). Yet linked biallelic variants can combine into multiple segregating haplotypes within short genomic regions, making genetic variation effectively *multiallelic* at the haplotype level. Although a complete description of linked variation requires recombination, e.g. in ancestral recombination graphs (Nielsen, Vaughn, and Deng, 2025), local haplotypes often provide a useful summary of short-range genetic diversity. Collapsing such variation into a biallelic framework may obscure important evolutionary dynamics and bias population genomic inference.

At a biallelic locus in a diploid population there are two possible homozygous states and a single heterozygous state. By contrast, a locus with *n* alleles can harbour up to *n* distinct homozygotes and *n*(*n* − 1)*/*2 distinct heterozygote genotypes (Gillespie, 2004; Hartl and Clark, 1997). When *n >* 2, multiple heterozygote types emerge, and once *n >* 3, heterozygotes outnumber homozygotes. Multiallelic loci therefore generate a qualitatively richer spectrum of genotypic diversity than biallelic loci, with potentially important consequences for how selection and drift shape allele frequency trajectories. We use the term *allele* in a *broad population-genetic sense* to denote a variant of the DNA sequence at a genomic region under consideration whose frequency can be tracked over the timescale of interest. Thus, depending on the application, an allele may correspond to a nucleotide variant at a single site, an alternative variant of a whole gene, a structural variant, or a local haplotype/founder-ancestry state within a genomic window.

Genetic diversity arises ultimately through mutation and is often elevated in genomic regions with high mutation rates — mutational hotspots — where multiple alleles may segregate simultaneously (Nesta, Tafur, and Beck, 2021). In addition, diversity may be actively maintained by balancing selection, including mechanisms such as heterozygote advantage and frequency-dependent selection (Hedrick, 2012; Siljestam and Rueffler, 2024; Ayala and Campbell, 1974; Ruzicka et al., 2025). Taken together, these considerations suggest that loci with more than two alleles should be common in natural populations, and that their evolutionary dynamics cannot, in general, be reduced to a biallelic approximation.

Empirically, multiallelic variation is widespread at biologically and medically important loci. The amylase gene, which is involved in starch digestion, exhibits extensive copy number variation in dogs, pigs, geladas, and humans (Axelsson et al., 2013; Ollivier et al., 2016; Paudel et al., 2013; Caldon et al., 2024; Perry et al., 2007). Other prominent examples, sometimes comprising hundreds of alleles, include the major histocompatibility complex (MHC), which plays a key role in vertebrate immunity (Radwan et al., 2020), the ABO blood group system and its many subgroups (Yamamoto, 2021), and genes underlying monogenic diseases such as cystic fibrosis (Sharma and Cutting, 2020). High allelic richness is also expected at loci involved in host–parasite interactions, where coevolutionary dynamics can maintain multiple alleles through balancing selection (Ebert and Fields, 2020; Cornetti et al., 2024). Similarly, self-incompatibility loci in flowering plants are known to exhibit extreme allelic diversity, thought to be driven by strong negative frequency-dependent selection (Takayama and Isogai, 2005; Wright, 1939; Yokoyama and Nei, 1979; Charlesworth, Vekemans, et al., 2005).

Recent advances in genome sequencing and analysis have made the empirical study of multial-lelic loci increasingly feasible. Detecting alleles segregating at low frequencies requires deep sequencing coverage (Gossmann and Waxman, 2022), which can be achieved either through pooled sequencing at high depth or by sequencing large numbers of individuals. While such approaches were previously prohibitively expensive, declining sequencing costs now allow rare alleles to be detected with increasing reliability. As a result, many loci previously classified as biallelic are likely to be revealed as multiallelic under deeper sampling. However, many commonly used summary statistics, such as Watterson’s *θ* and Tajima’s *D*, are primarily formulated for biallelic loci (Watterson, 1975; Tajima, 1989), and many analytical pipelines assume two alleles by default (Campbell et al., 2016). With the rapid growth in data richness there is a growing need for practical methodologies that can readily accommodate more than two alleles.

Here, we present a estimation procedure, based on the mathematical work of Shahshahani (1979) and Akin (1990), for the deterministic force of selection acting at a single locus. This allows a representation that has the desirable feature of explicitly disentangling the contribution of genetic diversity from that of genotype-specific fitness-effects, a feature that we shall exploit. This involves going beyond the results of Akin (1990), by expressing the selective force in terms of a matrix of fitness-effects of different genotypes. This matrix is analogous to the set of selection coefficients associated with different alleles. However, we do not assume genotype fitnesses are determined by additive effects of the alleles underlying a genotype. The situation is generally more complex than an additive scheme of selection. Our approach constitutes a practical computational approach that can accommodate an arbitrary number of alleles, a wide range of genotypic fitness schemes (including additive, multiplicative, fluctuating and frequency-dependent selection, along with heterozygote advantage). Using the *broad population genetic sense* definition of *allele*, as given above, this methodology can also be applied to tightly linked sets of sites, or equivalently to haplotypes, which is particularly relevant because selected haplotype blocks can persist over microevolutionary timescales and be reconstructed from replicated evolve-and-resequence time series (Franssen, Barton, and Schlötterer, 2017). When there are *n* alleles, the matrix of fitness-effects of genotypes contains *n*(*n* +1)*/*2−1 independent genotypic fitnesses, and this matrix, as a single entity, is the focus of our explorations.

In what follows, we show how to analyse allele frequency time series and to estimate genotype-level fitness-effect matrices over microevolutionary timescales (tens to hundreds of generations), such as those typical of experimental evolution studies. While this inference component is not intended as a fully general statistical solution, it is practical and accessible, despite the proliferation of genotypes associated with multiple alleles, and illustrates how multiallelic fitness interactions can be investigated using temporal data. Our results provide a practical foundation for the analysis of selection at multiallelic loci and for developing more comprehensive statistical extensions, e.g. fully likelihood-based or Bayesian inference methods.

## Motivation from a simple model

A very basic form of diversity is the genetic variation which exists at a single locus, and in what follows we shall often use *diversity* and *variation* as synonymous terms. Generally, natural selection acts on (biallelic or multiallelic) variation within a population and generates changes within the population. In addition to this, random genetic drift samples amongst existing variation within a population, and produces random changes. Thus both natural selection and random genetic drift only operate when there is variation within a population. This basic observation suggests that there may be a connection between the mathematical representation of these two distinct evolutionary ‘forces’ - selection and drift, which both vanish in the absence of variation. In this work we derive this connection by elementary considerations by co-opting an expression involving population variability, that appears in formulae for random genetic drift, to obtain a compact formula for the selective force when there are multiple alleles.

For the simplest model involving a locus with two alleles, the connection between selection and drift can be readily seen, as follows.

Consider an ‘ideal’ diploid sexual population (e.g., randomly mating hermaphrodites) with discrete generations, where a single biallelic locus is subject to weak additive or multiplicative selection. Let *x* denote the frequency of a focal allele at the locus in a given generation. Then by the beginning of the next generation we have that:

i. selection contributes the amount *sx*(1 − *x*) to the change in frequency, where *s* is a selection coefficient associated with the focal allele,
ii. random genetic drift contributes an amount *x*(1 − *x*)*/*(2*N*) to the variance of the frequency, where *N* is the census population size^1^,

(Gillespie, 2004). We note that these effects, from selection and drift, both contain an *identical function of frequency*, namely *x*(1 − *x*), which vanishes in the absence of variation (at *x* = 0 and *x* = 1). This shows a common dependency of these evolutionary forces on a basic measure of diversity of the population at the locus, namely *x*(1 − *x*).

In the mathematical literature it has been established that this observation, for two alleles, generalises to multiple alleles. That is, when there are more than two alleles at a locus, the effects of selection and random genetic drift have a common dependency on the diversity of the population (Akin, 1990).

When there are more than two alleles, what constitutes diversity is more complex than the case of two alleles; diversity is measured by a *matrix*. We shall accessibly motivate a variant of the mathematical result of Akin (1990), starting next, with a statement of the dynamics for a model involving selection that acts on multiple genotypes (produced by multiple alleles), and random genetic drift.

## Model

We adopt a Wright-Fisher model (Fisher, 1922; Wright, 1931) for a population of diploid hermaphroditic organisms, where generations are discrete and labelled by *t* = 0, 1, 2, At the start of each generation there are *N* adult organisms in the population. The quantity *N* is the *census size* of the population. The life cycle is as follows.

1. Reproduction A generation starts with *N* adults that randomly pair and produce offspring.
2. Adult death All adults die, with offspring remaining in the population.
3. Viability selection Offspring are subject to viability selection, which acts at a single locus with *n* alleles. The survivors of viability selection are termed juveniles.
4. Thinning Juveniles are randomly thinned to *N* individuals that become the adults of the next generation.

There is no restriction on the number of alleles at the locus, so that *n* can have the values 2, 3, 4, …, and we label the *j*’th allele as *A*_*j*_, with *j* = 1, 2, …, *n*.

To describe the population, we use an *n* component column vector **X**(*t*) whose *j*’th element, *X*_*j*_(*t*), is the frequency of allele *A*_*j*_ in adults, at the start of generation *t*, immediately prior to reproduction.

Rather than taking the *A*_*i*_*A*_*j*_ genotype to have a fitness proportional to *W*_*i,j*_ (relative fitness), we take the fitness to be proportional to 1 + *F*_*i,j*_, i.e.,

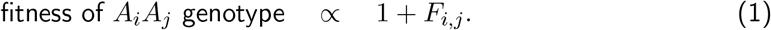

The quantity we introduce, *F*_*i,j*_, is a measure of the selection acting on the *A*_*i*_*A*_*j*_ genotype, and is analogous to a selection coefficient associated with an allele. Selection on different genotypes may have a complex origin that is generally not additive in effect. We shall often describe *F*_*i,j*_ as the ‘fitness-effect’ of the *A*_*i*_*A*_*j*_ genotype. The *F*_*i,j*_ may depend on allele frequencies, in which case selection is *frequency-dependent*, and in generation *t* the *F*_*i,j*_ will depend on **X**(*t*). We take the *F*_*i,j*_ to be constants unless we explicitly state otherwise.

Evolutionary dynamics depends on ratios of fitnesses, not absolute values. This means only relative fitnesses have a significance, and these are defined up to a multiplicative constant. In Appendix A we show the corresponding property of the *F*_*i,j*_.

In what follows, we shall use the symbol **F** to denote the *n* × *n* matrix whose (*i, j*) element is *F*_*i,j*_. Assuming the absence of genomic imprinting or other epigenetic phenomena (Peters, 2014) we have *F*_*i,j*_ = *F*_*j,i*_. That is, **F** is a *symmetric* matrix.

### Notation

For convenience we present here the remainder of the notation that we will use in the rest of this work.

Generally, the vector of allele frequencies at any time *t*, namely **X**(*t*), is a random variable with the properties of being non-negative and normalised to unity in the sense that *X*_*j*_(*t*) ≥ 0 and 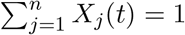. We write realisations (or non-random examples) of **X**(*t*) as **x** and these are also non-negative and normalised to unity.

With a T superscript denoting the *transpose* of a matrix:

- *δ*_*i,j*_ denotes a Kronecker delta, which takes the value 1 when the indices *i* and *j* are equal, and is zero otherwise
- **1** denotes an *n* component *column* vector with all elements 1:

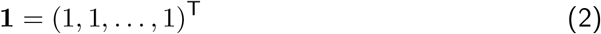
- ***I*** denotes the *n* × *n* identity matrix
- **V**(**x**) denotes an *n* × *n* matrix which depends on the vector **x**, and has elements given by

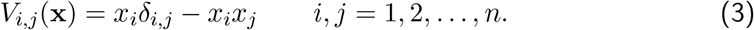

## Dynamics of the model

We first consider the behaviour of the population when its size is very large - effectively infinite. Let **X**(*t*) denote the column vector containing the *n* allele frequencies in the adults of generation *t*. The frequencies in generation *t* + 1 follow from the occurrence of reproduction and selection

in generation *t*, and are given by

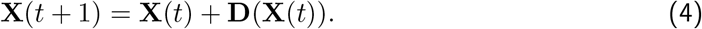

Here **D**(**X**(*t*)) is an *n* component column vector containing the deterministic changes that occur in allele frequencies in generation *t* due to natural selection. We will often call **D**(**X**(*t*)) the ‘selective force’ and will shortly give its form.

If, over the timescales of interest, we need also to take into account recurrent mutation between the different alleles, then we replace **X**(*t*) on the right hand side of Eq. (4) by **X**(*t*) + **MX**(*t*), where **M** is an *n* × *n* matrix containing mutation rates^2^.

When the population size is *finite*, in addition to selection, we also have the effects of a non selective thinning process, where *N* individuals are randomly picked to constitute the *N* adults of the next generation, with *N* denoting the census population size. The process of thinning introduces randomness, i.e., *random genetic drift*, into the dynamics, where the allele frequencies in generation *t* + 1 are distributed *around* the deterministic prediction given by the right hand side of Eq. (4). Incorporation of random genetic drift entails modifying Eq. (4) to

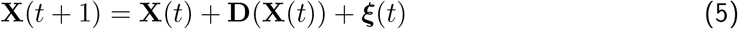

where ***ξ***(*t*) (also a column vector) is a random contribution to the change in allele frequencies.

To describe Eq. (5) in greater detail, we begin with the ‘force of genetic drift’, namely the vector ***ξ***(*t*), with elements *ξ*_*j*_(*t*). For any realisation of **X**(*t*), the expected value of ***ξ***(*t*) over replicate populations vanishes. We write this as *E* [***ξ***(*t*)|**X**(*t*) = **x**] = **0**, signalling that genetic drift is unbiased.

The variance-covariance matrix of ***ξ***(*t*), conditional on **X**(*t*) = **x**, is well known and given by^3^

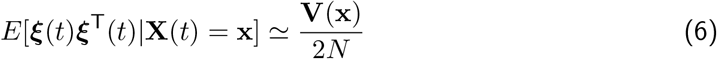

where the matrix **V**(**x**) of Eq. (3) enters - see Appendix B for details, and, as above, *N* denotes the census population size. Thus the (*i, j*) element of the variance-covariance matrix of ***ξ***(*t*) is given by *E*[*ξ*_*i*_(*t*)*ξ*_*j*_(*t*)|**X**(*t*) = **x**] ≃ *V*_*i,j*_(**x**)*/*(2*N*).

Consider now the selective force in Eq. (4), namely **D**(**X**(*t*)). There are different ways to write **D**(**x**) but in Appendix C we show that it can be compactly written as

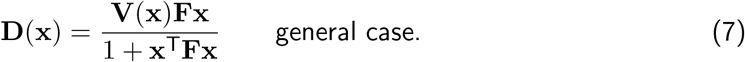

In this representation of **D**(**x**), which is equivalent to the force of selection in Akin (1990), the matrix **F** enters and is associated with genotypic fitness - see Eq. (1). But we also see the matrix **V**(**x**), which naturally emerged as a feature of random genetic drift. Evidently, the matrix **V**(**x**) also plays a key role in the form taken by the evolutionary force of selection^4^

The connection between selection and drift is even clearer under the continuous time, continuous state, *diffusion approximation* of Eq. (5) when selection is weak, in the sense that all elements of **F** are small (|*F*_*i,j*_| ≪ 1). In this case we can neglect the **F** dependence in the denominator of Eq. (7), with the approximation^5^ **D**(**x**) ≃ **V**(**x**)**Fx**. The diffusion approximation then says over a small time interval *dt*, the change in frequencies, *d***X**, arises from the sum of effects of selective and drift forces. The diffusion approximation of Eq. (5) is motivated in Appendix B and omitting *t* arguments it takes the form

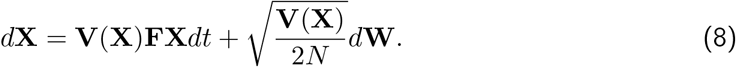

Here **W** ≡ **W**(*t*) is a column vector containing random elements, representing the stochasticity of random genetic drift^6^.

We see in Eq. (8) that precisely the same frequency-dependent matrix, **V**(**X**), appears in the term arising from selection, **V**(**X**)**FX***dt*, and the term arising from random genetic drift, 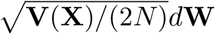. We associate this matrix with the *diversity* of the population. In particular, the *i*’th diagonal element of **V**(**X**), namely *X*_*i*_(1 − *X*_*i*_) is clearly recognisable as a measure of diversity. Furthermore, when a single allele is at a frequency of unity (and all other alleles at zero frequency), all elements of **V**(**X**) vanish. However, the matrix **V**(**X**) is also associated with a notion of *directionality*. The matrix **V**(**X**) plays an instrumental role in allele frequency *changes* that originate in selection or drift. These changes take place in the space of the *n* allele frequencies, along a direction that is dependent on properties of the matrix **V**(**X**), namely its eigenvectors.

Overall, the matrix **V**(**X**) is somewhat richer than just a direct measure of diversity, and while it contains a measure of diversity, and vanishes in the absence of diversity, it also ‘directs’ allele frequency changes. In multiallelic problems we will, for convenience, just refer to **V**(**X**) as the *diversity*.

With Eqs. (7) and Eq. (8), we have motivated, in relatively elementary terms, the results present in the mathematical literature (Shahshahani, 1979; Akin, 1990) that when there are multiple alleles, selection depends on the *diversity* of the population, as does random genetic drift. Furthermore, when allele frequencies are given by **x**, diversity is described by the matrix **V**(**x**).

## Special cases of D(**x**)

We next explore some special cases of the selective force given in Eq. (7).

### Additive selection

Under additive selection, the *A*_*i*_*A*_*j*_ genotype has a fitness proportional to 1 + *F*_*i,j*_ = 1 + *s*_*i*_ + *s*_*j*_ where *s*_*i*_ is a selection coefficient associated with allele *A*_*i*_. With **s** an *n* component column vector whose *i*’th element is **s**_*i*_, we have

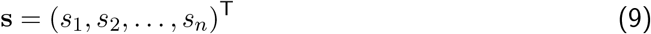

and under additive selection we can write the matrix **F** as

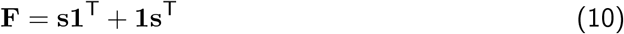

which yields *F*_*i,j*_ = *s*_*i*_ + *s*_*j*_.

Given that **x** represents a column vector of possible allele frequencies, it has the property 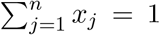, which can be written as **1**^T^**x** ≡ **x**^T^**1** = 1, and this leads to **V**(**x**)**1** vanishing (i.e., using Eq. (3) it becomes a vector with all elements zero). Thus **V**(**x**)**Fx** = **V**(**x**) (**s1**^T^ + **1s**^T^**)x** = **V**(**x**)**s** and also 1 + **x**^T^**Fx** = 1 + **x**^T^ (**s1**^T^ + **1s**^T^**)x** = 1 + 2**s**^T^**x**. These lead to

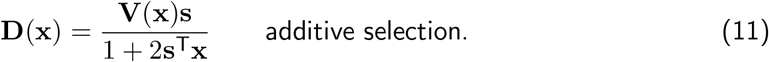

In the special case of *n* = 2 alleles, with *s*_1_ = *s, s*_2_ = 0, and writing *x* = *x*_1_, we obtain a selective force on the frequency of allele *A*_1_ of *D*_1_(**x**) = *sx*(1 − *x*)*/* (1 + 2*sx*).

### Multiplicative selection

Under multiplicative selection, the *A*_*i*_*A*_*j*_ genotype has a fitness proportional to 1 + *F*_*i,j*_ = (1 + *s*_*i*_) (1 + *s*_*j*_) and again we shall refer to *s*_*i*_ as the selection coefficient associated with allele *A*_*i*_. In this case, again using Eq. (9), we can write

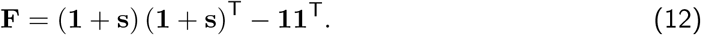

We then have **V**(**x**)**Fx** = **V**(**x**)**s (**1 + **s**^T^**x)** and 1 + **x**^T^**Fx** = (1 + **s**^T^**x)**^2^ and hence

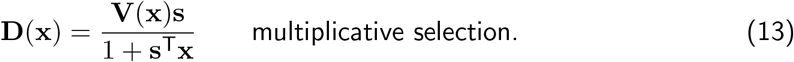

In the special case of *n* = 2 alleles, with *s*_1_ = *s, s*_2_ = 0, and writing *x* = *x*_1_, we obtain a selective force on the frequency of allele *A*_1_ of *D*_1_(**x**) = *sx*(1 − *x*)*/* (1 + *sx*).

### Heterozygote advantage

We next consider one of the simplest forms of selection that acts genuinely at the level of genotypes, and cannot be devolved to properties of individual alleles. This is selection where all heterozygotes have a common fitness advantage of *σ* relative to all homozygotes. That is

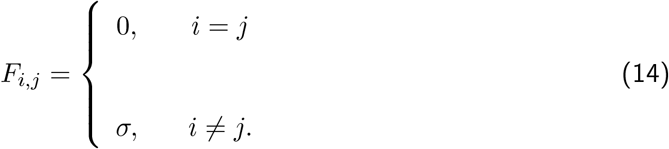

This corresponds to

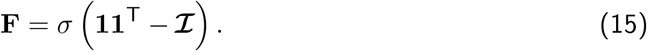

Using this result in Eq. (7) yields

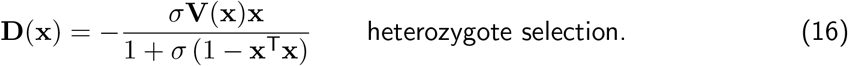

In the special case of *n* = 2 alleles, and writing *x* = *x*_1_, we obtain a selective force on the frequency of allele *A*_1_ of *D*_1_(**x**) = *σx*(1 − *x*)(1 − 2*x*)*/* [1 + 2*σx*(1 − *x*)].

### Fluctuating selection

We consider the situation where fitnesses randomly change (Huerta-Sanchez, Durrett, and Bustamante, 2008; Gossmann, Waxman, and Eyre-Walker, 2014), from one generation to the next, due, for example, to random environmental effects on the population. Here we give one example based on Eq. (7).

Assuming fitnesses are not correlated over time, it is then appropriate to directly average over the fluctuating fitnesses. The standard way of proceeding, *as far as the selective force is concerned* (we do not consider the change in the variance of allelic effects), is to first expand the selective force to second order in selection coefficients, and then average the result (Jensen and Pollak, 1969). Here, we shall interpret the analogue of selection coefficients as elements of the matrix **F**, that appears in Eq. (7), and we expand **D**(**x**) to second order in **F**. This leads to **D**(**x**) ≃ **V**(**x**)**Fx** − **V**(**x**)**Fx (x**^T^**Fx)**.

As an illustrative example, we take

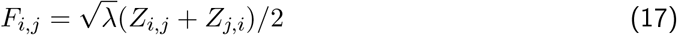

where the *Z*_*i,j*_ are, for all *i* and *j*, independent normal random variables with mean zero and variance one, while *λ* is a measure of variance of fitness-effects.

The form of *F*_*i,j*_ adopted in Eq. (17) ensures that *F*_*i,j*_ = *F*_*j,i*_. It also leads to the mean value of **F** vanishing (the average effect of selection on any genotype is zero) which we write as 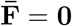. Furthermore with *F*_*i,j*_ given by Eq. (17), to calculate the average of **D**(**x**), we require knowledge of the average of *F*_*i,j*_*F*_*k,l*_ for arbitrary *i, j, k* and *l*. This is given by

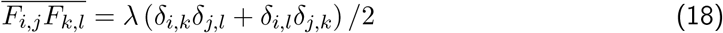

and using this result leads to^7^

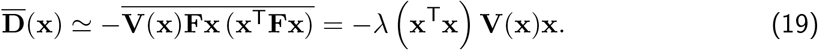

In the special case of *n* = 2 alleles, and writing *x* = *x*_1_, we obtain an averaged selective force on allele *A*_1_ of 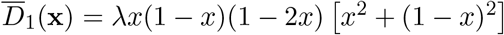.

### Frequency-dependent selection

We can model negative frequency-dependent selection by allowing the fitness of a genotype to depend on the genotype’s frequency, such that the fitness is a *decreasing function* of genotype frequency. This results in the matrix of fitness-effects, **F**, now becoming **F**(**x**), where **x** can be thought of as the vector of current allele frequencies. The *A*_*i*_*A*_*j*_ genotype now has a fitness proportional to 1 + *F*_*i,j*_(**x**). Given that on production, this ordered genotype has a frequency of *x*_*i*_*x*_*j*_, we take

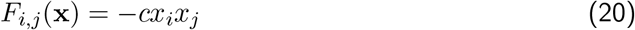

where *c* is a constant^8^ that lies in the range 0 ≤ *c <* 1.

Since we can write **F**(**x**) = − *c* **xx**^T^, the form of the selective force following from Eq. (7) is

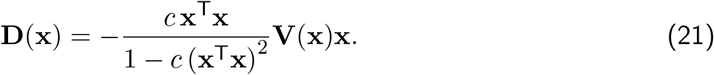

In the special case of *n* = 2 alleles, and writing *x* = *x*_1_, we obtain a selective force on the frequency of allele *A*_1_ of 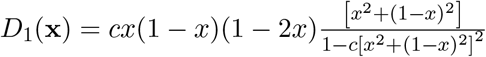.

### Illustrations of the dynamics

To illustrate some interesting dynamics, we calculated allele frequency trajectories numerically in MATLAB (The Mathworks Inc., 2025) in the absence of mutation. The detailed scripts are available as a github repository (https://github.com/NikolasVellnow/selection_multiallelic). Purely for illustrative purposes, we present results for *n* = 3 alleles; the representation of selection presented can apply for an arbitrary number of alleles, and indeed later we consider empirical data for a case with *n* = 4 alleles. Thus for a locus with *n* = 3 alleles, we have calculated frequency trajectories for a number of different scenarios, over 5 × 10^3^ generations. In each scenario we calculated three stochastic trajectories, corresponding to a (census) population size of 10^3^ individuals, and a large number of deterministic trajectories, corresponding to an effectively infinite population.

In the results we present, the finite (but large) number of generations we consider has the consequence that the deterministic trajectories closely approach, but never actually reach the end point they would ultimately achieve. In what follows, we shall make statements about ‘achieving fixation’ or ‘achieving a stable polymorphism’ that should be interpreted to hold to an accuracy of machine precision (1 part in 10^14^).

#### Trajectories for a single randomly generated set of fitness-effects

In Figure 1, we proceeded by first calculating the dynamics with a randomly generated set of fitness-effects, based on a normal distribution, as given in Eq. (17). We generated the set of fitness values, corresponding to the resulting **F** in Eq. (22), only once at the start of the simulation. We then used Eq. (7), which applies for the general case, to calculate the allele frequency dynamics.

**Figure 1.**
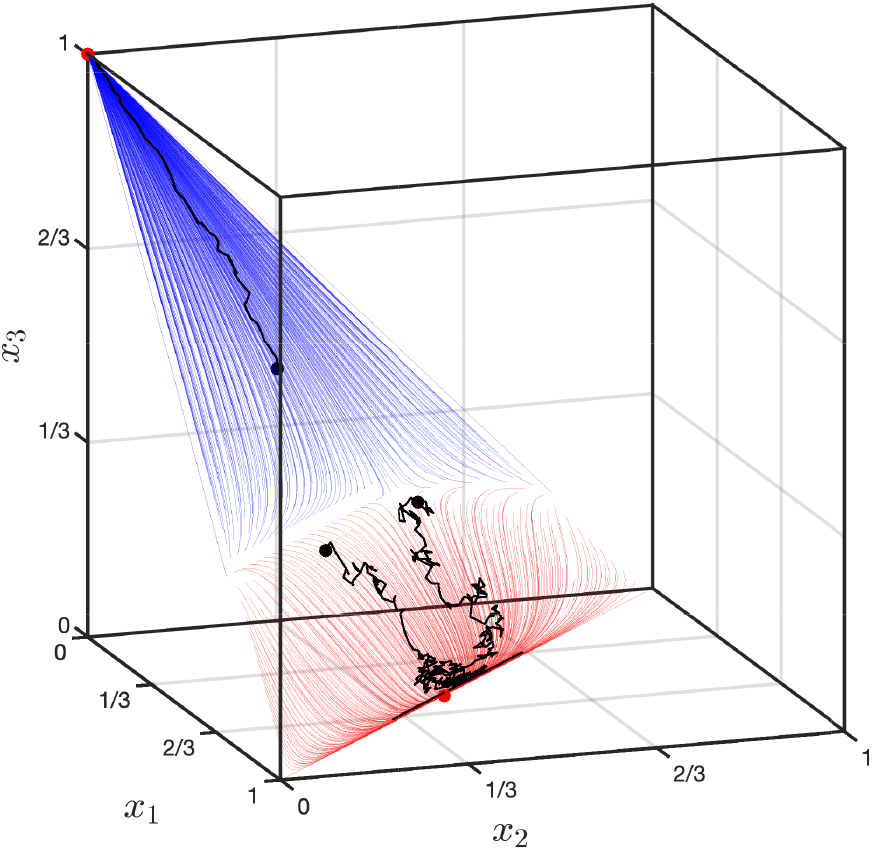
Plots of allele frequency trajectories for *n* = 3 alleles, for a single randomly generated set of fitness-effects. In the figure, allele frequencies are denoted by *x*_1_, *x*_2_ and *x*_3_. The figure contains three stochastic trajectories, which are plotted in black. These start from distinct points (allele frequency sets) indicated by the three black dots. The population size associated with these stochastic trajectories is *N* = 10^3^. In addition, a large number of deterministic trajectories, associated with an effectively infinite population, are plotted. These have end points indicated by the two red dots. The deterministic trajectories arise from many different initial points and are coloured according to the final point they reach. Both stochastic and deterministic trajectories were run for a total of 5 × 10^3^ generations.

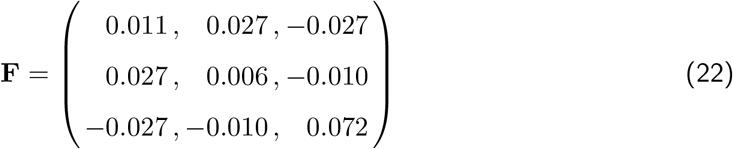

We simulated three stochastic trajectories, for the populations of finite size, that started from the following three sets of allele frequencies: (0.50, 0.25, 0.25), (0.25, 0.50, 0.25), and (0.25, 0.25, 0.50). We also produced a large number of deterministic trajectories, for the populations of effectively infinite size, with a variety of different starting points. We observe that the end points of the deterministic trajectories depend on the initial frequencies (see Figure 1). Deterministic trajectories with starting frequencies in the “lower area” all reached the end point (0.5595, 0.4405, 0) (see Figure 1; red trajectories). This end point, under deterministic dynamics, is a stable polymorphism, where alleles *A*_1_ and *A*_2_ are maintained in the population while allele *A*_3_ is lost. By contrast, deterministic trajectories with starting frequencies in the “upper area” reached the end point (0, 0, 1), corresponding to fixation of allele *A*_3_ (see Figure 1; blue trajectories).

Some theoretical results on polymorphic equilibria with multiple alleles can be obtained from the zero recombination limit of multilocus models (see e.g., Christiansen, 1999).

#### Trajectories for a single set of additive fitness-effects

In Figure 2 we illustrate the dynamics when fitnesses are additive (Eq. 10). For this, we simulated three stochastic trajectories that started from the following three sets of allele frequencies: (0.70, 0.15, 0.15), (0.15, 0.07, 0.15), and (0.15, 0.15, 0.70). The set of fitness values was calculated from the additive allelic selection coefficients (0.01, 0.03, 0.06) which were then used throughout. We also produced many deterministic trajectories (which all started with three alleles segregating). The stochastic and determinsitic trajectories were all observed to reach fixation of the *A*_3_ allele (see Figure 2).

**Figure 2.**
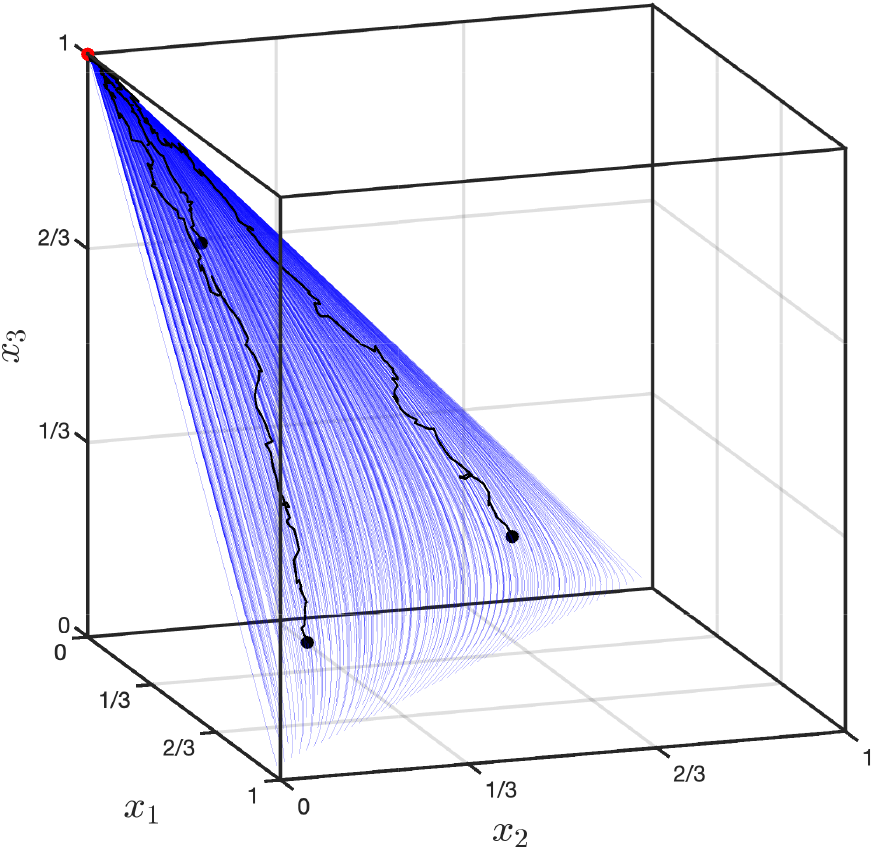
Plots of allele frequency trajectories for *n* = 3 alleles, for a single set of additive fitness-effects. In the figure, allele frequencies are denoted by *x*_1_, *x*_2_ and *x*_3_. The figure contains three stochastic trajectories, which are plotted in black. These start from distinct points (allele frequency sets) indicated by the three black dots. The population size associated with these stochastic trajectories is *N* = 10^3^. In addition, a large number of deterministic trajectories, associated with an effectively infinite population, are plotted. The deterministic trajectories arise from many different initial points and have a common end point, indicated by the red dot. All deterministic trajectories are coloured blue. Both stochastic and deterministic trajectories were run for a total of 5 × 10^3^ generations.

#### Trajectories for negative frequency-dependent selection

In Figure 3 we give an example of balancing selection, with the dynamics calculated under frequency-dependent selection (Eq. 20). We simulated three stochastic trajectories, that had the starting allele frequency sets (0.70, 0.15, 0.15), (0.15, 0.70, 0.15), and (0.15, 0.15, 0.70). The strength of frequency-dependent selection was *c* = 0.05. The set of fitness values was updated every generation, based on the current allele frequencies following Eq. (20)). Here, all deterministic trajectories reached the same stable polymorphism, that had the intermediate end point frequencies (1*/*3, 1*/*3, 1*/*3), independent of the starting frequencies (see Figure 3; blue trajectories). All populations following a stochastic trajectory drifted around the point (1*/*3, 1*/*3, 1*/*3) and, over the time of simulation, exhibited a polymorphism that included all alleles (see Figure 3)

**Figure 3.**
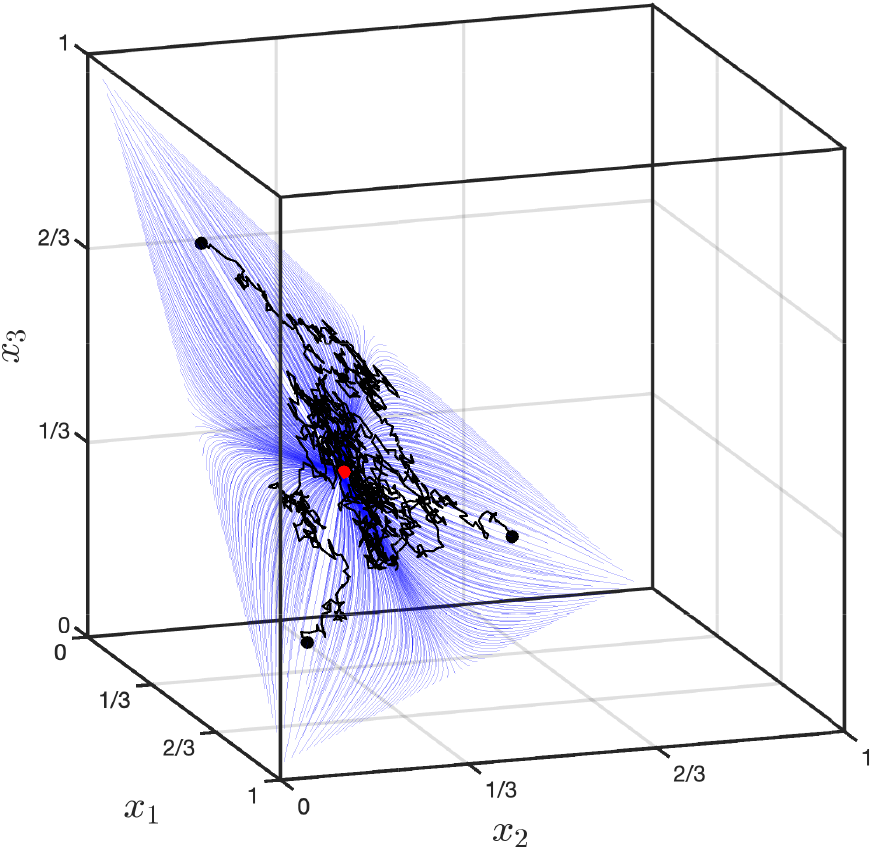
Plots of allele frequency trajectories for *n* = 3 alleles, for negative frequency-dependent selection. In the figure, allele frequencies are denoted by *x*_1_, *x*_2_ and *x*_3_, and a single form of fitness-effects, associated with negative frequency-dependent selection, was used. The figure contains three stochastic trajectories, which are plotted in black. These start from distinct points (allele frequency sets) indicated by the three black dots. The population size associated with these stochastic trajectories is *N* = 10^3^. In addition, a large number of deterministic trajectories, associated with an effectively infinite population, are plotted. The deterministic trajectories arise from many different initial points and have a common end point, indicated by the red dot. All deterministic trajectories are coloured blue. Both stochastic and deterministic trajectories were run for a total of 5 × 10^3^ generations.

#### Incorporating diverse cases into the model

To generate evolutionary trajectories for other diverse cases of the selective force, it is only necessary to change the form of the matrix **F**(**x**), and decide if it is calculated only once (e.g., multiplicative selection), at the beginning of the simulation, or whether it has to be updated after each generation (e.g., frequency-dependent selection). This highlights the flexibility with which our approach can implement different forms of selection on multiallelic loci.

## Applications to empirical data analysis

Our approach can also be useful in understanding and analysing empirical data. In general, it can be applied to all population genomic data sets that allow the tracking of multiple allele or haplotype frequencies over several generations. For instance, evolve-and-resequence experiments are prime candidates to be analysed in this way (Schlötterer et al., 2015; Franssen, Barton, and Schlötterer, 2017).

### Estimating F from allele frequency trajectories

One application of this work is the estimation of **F** from empirical trajectories of multiple alleles or haplotypes. We obtain a regularised point estimate of **F** that summarises the allele-frequency changes under the assumed model. This estimate provides an interpretable representation of the fitness interactions associated with the observed trajectories and can be used to compare genotypes, identify candidate heterozygote advantage, and guide further analyses. Note that estimation of **F** is not possible from allele frequencies at one or two time points. With *n* alleles, the matrix **F** has a total of *n*(*n* + 1)*/*2 − 1 independent elements, while there are only *n* − 1 independent allele frequencies. Thus to estimate **F** we need to consider allele frequency *trajectories*. We shall thus estimate **F** from empirical *trajectories* of multiple alleles or haplotypes. The present implementation focuses on point estimation; extensions incorporating formal uncertainty quantification, explicit sampling models, and model comparison would provide valuable further developments.

Here, we present an algorithm to estimate the matrix of fitness-effects, **F**, when the following assumptions are applicable:

i. the **F** matrix is constant (i.e., has no time or frequency dependence), and
ii. over the time interval considered, mutations can be neglected.

Under these assumptions, the equation that determines allele frequency trajectories is Eq. (5).

Then with a data set consisting solely of empirical allele frequency trajectories, that cover a range of times, we proceed as follows.

First, we define a ‘cost function’ *C*(**F**) which measures the mismatch between the empirical allele frequency trajectories of the data set, and the allele frequency trajectories arising from a putative matrix of fitness-effects, **F**. The cost function incorporates the Shahshahani metric (Shahshahani, 1979; Akin, 1990). This leads to the cost being proportional to a large *N* approximation of log likelihood (Akin, 1990). Then, using a suitable optimization algorithm we find the matrix of fitness-effects that minimizes the cost function, *C*(**F**). The estimated matrix of fitness-effects produces trajectories that are similar to the empirical trajectories. Details of the estimation algorithm are given in Appendix D.1, with additional notes given in Section 1 of the Supplementary Material. Since, in real world data collections, genomic data is often not available for every generation, but instead may be given for time points with different numbers of generations between them, we also provide an algorithm that can estimate the matrix of fitness-effects in such cases - see Appendix D.2.

### Application to a yeast data set

As an example of estimating the matrix of fitness-effects, on real world data, we analysed publicly available data from an evolve-and-resequence experiment with the outcrossing brewer’s yeast *Saccharomyces cerevisiae* (Burke, Liti, and Long, 2014; Phillips et al., 2020). Briefly, the authors crossed four wild yeast strains to create a mixed outbred founder population and let replicated populations evolve for 540 generations under a laboratory domestication treatment. Sexual reproduction was enforced every 30 generations and samples of the populations were resequenced at 17 time points during the experiment. The authors provided the relative frequencies of haplotypes from the four original wild strains, calculated for 10 KB windows across 17 time points, enabling the tracking of their evolutionary trajectories (Burke, Liti, and Long, 2014; Phillips et al., 2020). Here, we take the four haplotypes as being equivalent to four alleles at a locus in a diploid organism, which is consistent with previous approaches in which selected haplotype blocks are inferred from correlated allele-frequency changes over time (Franssen, Barton, and Schlötterer, 2017). Since the four founder haplotypes observed within a 10 KB window are treated as four local allelic states the fitted matrix therefore describes effective founder-haplotype fitness interactions for that genomic region, integrating the effects of linked variation that shape the local haplotype trajectories. It should not be interpreted as identifying the causal fitness-effects of individual variants within the window. Furthermore, all 30 generations of mitotic reproduction, between outcrossing events, are taken as a single effective generation of a diploid organism, during which time selection occurs.

First, as an example region, we estimated **F** (with minimum cost of 0.620) for the trajectories of the haplotypes in the 10 KB window around position 457,847 on chromosome C16 for replicate population 3 (see Figure 4a; cf. Figure 6b in Phillips et al., 2020). The estimated **F** has a form which depends on which genotype is taken as the reference type (other, ‘evolutionarily equivalent’ forms of **F**, which use different reference genotypes, can be obtained from the transformation in Appendix A). Throughout, we label the four founder haplotypes as *A*_1_ = DBVPG6044, *A*_2_ = DBVPG6765, *A*_3_ = Y12, and *A*_4_ = YPS128.

**Figure 4.**
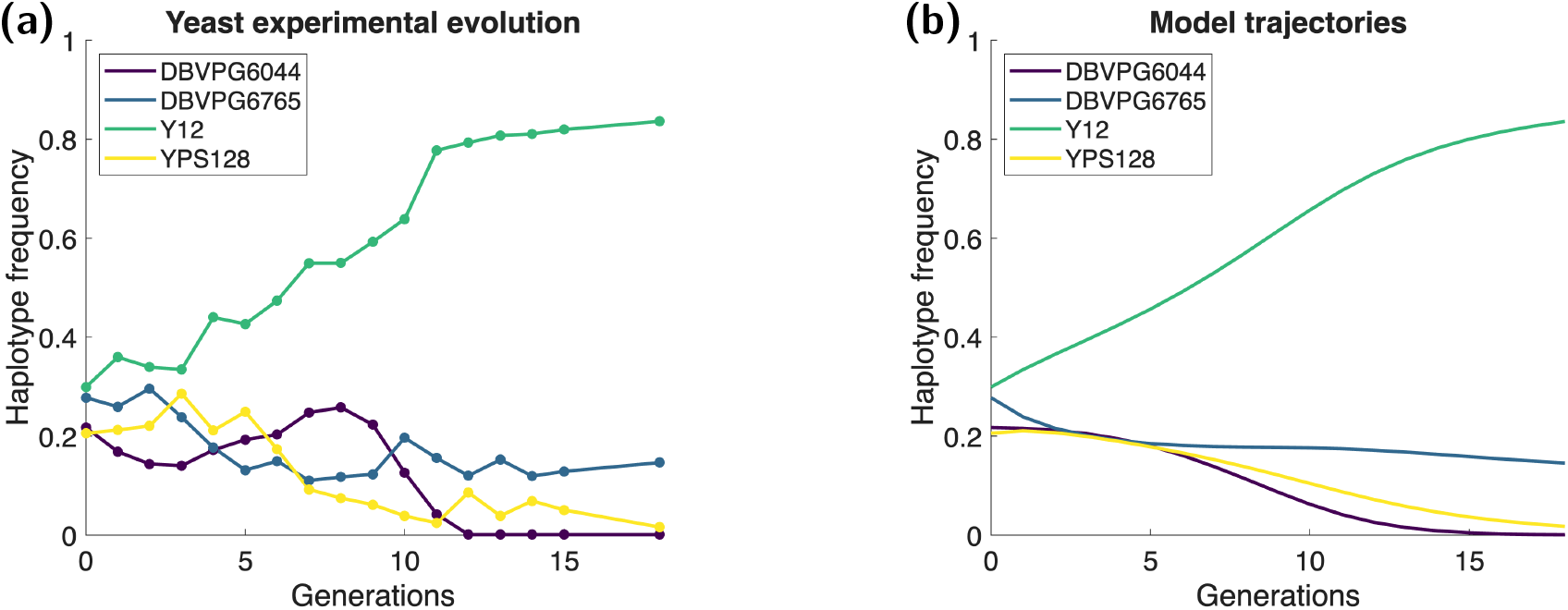
Comparison of haplotype frequency trajectories from a yeast experiment and from estimated parameters using our new model. Data is from a 10 KB window around position 457,847 on chromosome C16 for replicate 3 (Burke, Liti, and Long, 2014; Phillips et al., 2020). Haplotypes are DBVPG6044, West African in dark blue, DBVPG6765, wine/European in purple, Y12, sake/Asian in green, and YPS128, North American in yellow. a) Empirical trajectories from yeast experimental evolution. b) Trajectories produced by Eq. (4) where the estimated **F** matrix, Eq. (23), was used.

Here, we give the form of **F** when the genotype *A*_3_*A*_4_ is used as the reference type (in which case its fitness-effect vanishes). We also give the corresponding matrix of relative fitness values, which we denote by **w**. This matrix contains the fitness values of all genotypes, *relative* to that of the most fit genotype, i.e., *w*_*i,j*_ = (1 + *F*_*i,j*_)*/*(1 + max_*k,l*_(*F*_*k,l*_)):

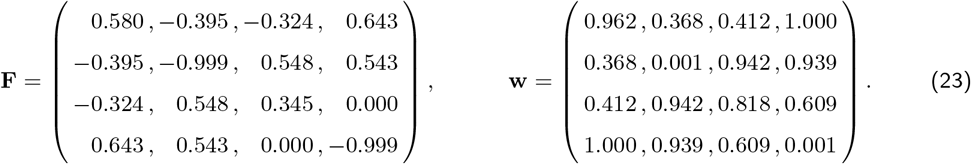

Using the above form of **F** produces trajectories that closely capture the main trends of the empirical haplotype-frequency data (see Figure 4).

Once **F**, and hence **w**, have been estimated from the data, they can be used to address questions of interest. For instance, within this fitted effective model, heterozygote genotypes have, on average, a relative fitness approximately 60% higher than homozygote genotypes in this region. However, the individual heterozygote combinations are heterogeneous: some show pronounced advantages (*A*_1_*A*_4_, *A*_2_*A*_3_, and *A*_2_*A*_4_), whereas others are less favourable than the corresponding homozygotes (*A*_1_*A*_2_ and *A*_1_*A*_3_). Because adjacent windows retain linkage, these results are most naturally interpreted as a genome-wide pattern of effective founder-haplotype interactions rather than as independent estimates of selection at individual 10 kb loci.

The results we obtain, for the estimate of **F**, are sensitive to the validity of the assumptions we have made. When applying our estimation method to regions of another chromosome (chromosome C12), we do not see such a good fit to the data (see Figure S1). We suspect that one or more of the assumptions we have made may not apply. However, the effective population size of *N*_*e*_ ≈ 10^6^ quoted for this experiment (Burke, Liti, and Long, 2014) does not suggest a dominant role for genetic drift, although we discuss caveats regarding this estimate in the Supplementary Material (Section 4).

We next fitted effective founder-haplotype fitness matrices across 1,161 non-overlapping 10 KB windows spanning the 16 major chromosomes of *S. cerevisiae*. Across these fitted matrices, the highest relative-fitness entry was a heterozygote in 99.8 % of windows, with the *A*_3_*A*_4_ genotype being the most fit in 29.4 % of cases (see Figure 5), a pattern that a null-model comparison shows is highly unlikely to arise by genetic drift alone (see Supplementary Section 4 and Figures S4–S5). This widespread pattern highlights the capacity of the framework to reveal heterogeneity in inferred genotype-level fitness interactions across the genome. Because adjacent windows retain linkage, these results are most naturally interpreted as a genome-wide pattern of effective founder-haplotype interactions rather than as independent estimates of selection at individual 10 KB loci.

**Figure 5.**
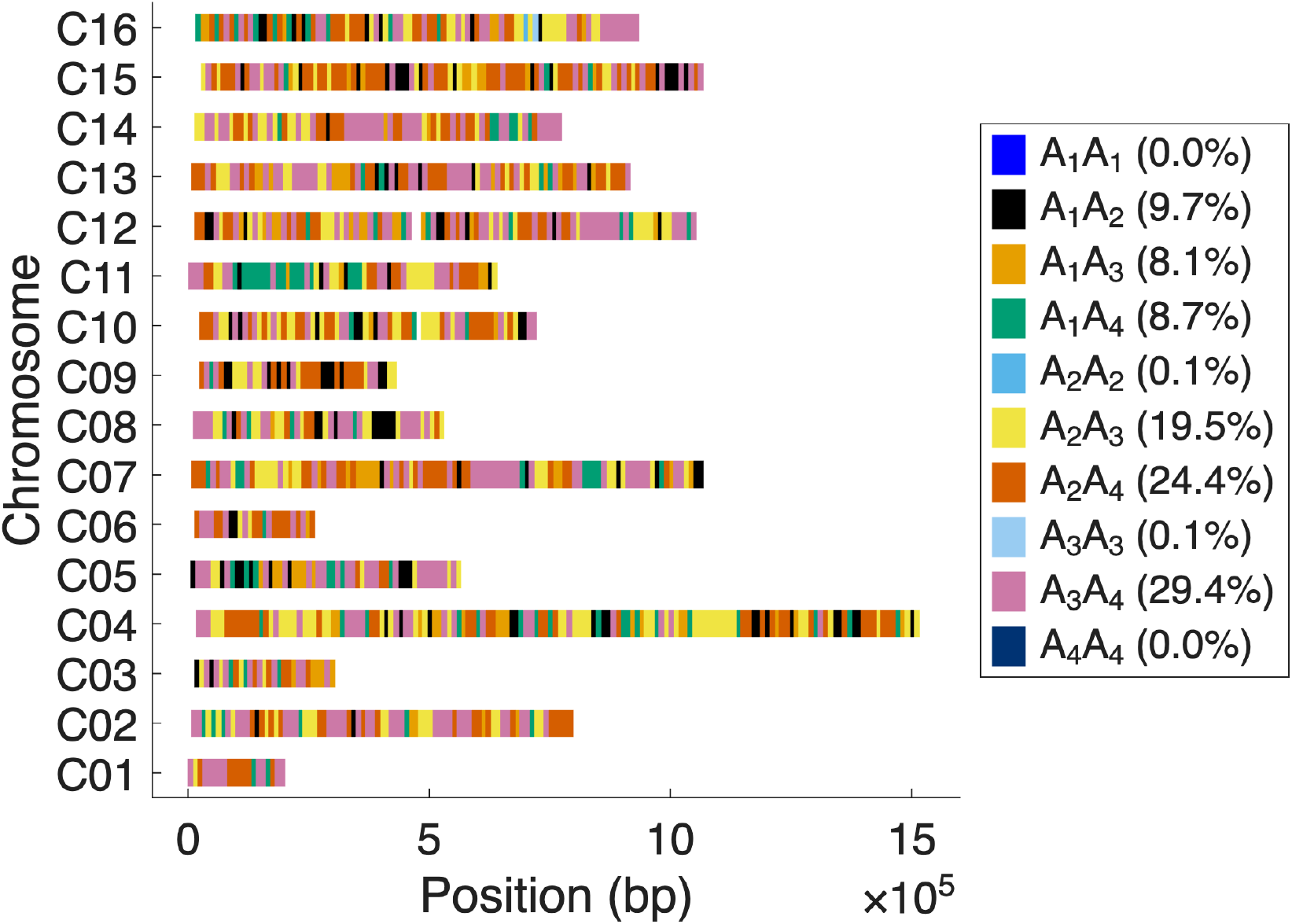
Illustration of the most fit genotypes across the major 16 S. cerevisiae chromosomes based on the experimental evolution experiment by Burke, Liti, and Long (2014) and Phillips et al. (2020). Each colored point indicates which genotype was the fittest for non-overlapping 10 KB windows (plotted at their mid-point), using the haplotype labelling *A*_1_ = DBVPG6044, *A*_2_ = DBVPG6765, *A*_3_ = Y12 and *A*_4_ = YPS128. Homozygotes are colored in shades of blue, while heterozygotes are colored in black, orange, bluish green, yellow, vermillion, and reddish purple. Relative frequencies with which each genotype was found to be the fittest one are given in brackets in the legend. Note that the data provided does not always start at the first possible mid-point (5 KB) for all chromosomes so that the plot starts at different positions on the left hand side.

## Discussion

In this work we have presented a general framework for multiallelic selection that represents population diversity and genotype-level fitness effects through the matrices **V**(**x**) and **F**, respectively. This representation provides a unified and flexible description of a broad range of selection regimes acting on an arbitrary number of alleles. We additionally demonstrated how such fitness-effect matrices can be determined from allele- or haplotype-frequency time series, thereby connecting this work to empirical evolutionary data. Our practical framework has its origins in previous mathematical results (Shahshahani, 1979; Akin, 1990) and can be used to investigate multiallelic genetic diversity that incorporates complex forms of selection. The fitting procedure presented provides a useful starting point for empirical applications and can naturally be extended to incorporate formal uncertainty estimates, and comparisons among alternative evolutionary models.

The method allows estimation of the selective force on different genotypes from empirical data in the presence of some degree of genetic drift. We have provided detailed scripts as supplemental material to this manuscript, which allow interested researchers to tailor our approach to their specific research questions. Note, in particular, that our approach allows the incorporation of complex forms of selection that act at the level of genotypes and, importantly, can be applied when there are an arbitrary number of alleles. Therefore, unlike treatments that assume biallelic loci, our approach can incorporate additional alleles that may be present at very low frequencies. This is particularly relevant: (i) when the genetic diversity of a single locus is very high and many alleles exist (Radwan et al., 2020; Yamamoto, 2021; Sharma and Cutting, 2020), or (ii) when the effective population size at a particular locus is very high (Gossmann, Woolfit, and Eyre-Walker, 2011; Smith and Haigh, 1974; Charlesworth, Morgan, and Charlesworth, 1993).

The inference approach presented here is intended primarily as a conceptual and methodological foundation rather than as a complete statistical solution. Inference of the fitness matrix **F** is undoubtedly complicated, and inherently sensitive to modelling choices. In addition, estimates can be influenced by the treatment of rare alleles, whose trajectories are particularly susceptible to stochastic sampling effects, and more generally, by noise in allele frequency estimates. These features are intrinsic to multiallelic time-series data and are not unique to the present approach. Addressing them, fully appropriately, will require extensions that incorporate explicit models of sampling error, the stochasticity of the dynamics, and formal measures of uncertainty. We view the framework developed here as a starting point for such future work, providing a unified representation of multiallelic selection upon which likelihood-based or Bayesian inference methods, model comparison, and robustness analyses can be built.

Several statistical methods have been developed to infer selection from allele-frequency time series, typically within Wright–Fisher or diffusion-based frameworks. Likelihood-based and Bayesian approaches have been used to estimate allele-specific selection coefficients from temporal data in both experimental and natural populations (Malaspinas, 2016; Whitehouse and Schrider, 2023; Garcia Castillo et al., 2024). Such approaches have proven powerful in analyses of evolve-and-resequence experiments and ancient DNA datasets, but generally focus on biallelic loci and estimate selection coefficients associated with individual alleles. In contrast, the framework developed here accommodates an arbitrary number of alleles and focuses on estimating genotype-level fitness interactions through the matrix of fitness-effects **F**, thereby extending time-series inference beyond the commonly assumed biallelic setting.

Although the estimation framework developed here is based on deterministic allele-frequency dynamics, it naturally raises the question of how stochastic effects should be incorporated into multiallelic evolutionary inference. Our estimation procedure already incorporates such considerations, at least approximately: the Shahshahani metric underlying our cost function is proportional to a large-*N* approximation of the log-likelihood under multinomial sampling (Akin, 1990), so that our fitted **F** reflects, to leading order, the effects of genetic drift rather than assuming a purely deterministic process. Because this metric weights allele-frequency deviations by 1*/X*_*i*_(*t*), it places particularly strong weight on rare alleles, where stochastic effects are most pronounced; we address this via the small constant *α* added to the denominator (Eq. 48), together with the ridge penalty that further stabilises estimates.

A natural next step would be to replace this large-population approximation with a fully likelihood-based or Bayesian inference framework that explicitly models stochastic allele-frequency trajectories while simultaneously providing formal uncertainty estimates for the inferred fitness matrix **F**.

More broadly, rare alleles remain particularly susceptible to stochastic loss, and increasing allelic richness generally leads to increasingly complex allele-frequency dynamics. Over evolutionary timescales, genetic drift therefore acts to erode genetic diversity (Kimura, 1955; Latter and Novitski, 1969; Vellnow, Gossmann, and Waxman, 2024), whereas balancing selection, including heterozygote advantage and frequency-dependent selection, can counteract this process and maintain polymorphism (Kimura and Crow, 1964; Lewontin, Ginzburg, and Tuljapurkar, 1978; Spencer and Marks, 1988; Siljestam and Rueffler, 2024). Correctly identifying the full spectrum of allelic variation is therefore essential for understanding the interplay between mutation, drift, and balancing selection. Our framework provides a natural way to investigate such scenarios, including cases in which only specific allelic combinations give rise to heterosis.

Since our model allows the incorporation of complex forms of selection, which act at the level of the genotype, it can be used to study a wide range of questions. For this, it is only necessary to define the matrix of fitness-effects of relevance to the specific research question. For example, we believe the interesting model from Haasl and Payseur, where genotypic fitness depends on the length of a microsatellite allele at a specific locus (Haasl and Payseur, 2013; Haasl, Johnson, and Payseur, 2014), could be modelled as a special case of our matrix of fitness-effects. Furthermore, arbitrarily complex forms of incomplete dominance can also be included.

There are some direct spin-offs from the general framework that we have presented. For example, it exposes relations between the selective forces arising from different forms of fitness. Consider the selective forces in Eqs. (16), (19), and (21), that arise, respectively, from fitnesses that act on heterozygotes, or are fluctuating, or that lead to negative frequency-dependent selection. While all these forms of fitness are different, the selective forces that arise all have very similar forms. In particular all three expressions for the force are of the form −*κ***V**(**x**)**x**, where *κ* depends on **x** but has a purely numerical value (it is not a vector or matrix). This means all three selective forces push allele frequencies in precisely the same ‘direction’, but by different amounts, because the coefficient, *κ*, generally has different forms and different values in the three cases.

In our description of the dynamics of the model, we mentioned the way recurrent mutation is incorporated into the evolutionary forces that generate allele frequency changes. There is another issue associated with mutation that we should mention. Consider the situation where we have allele frequency trajectory data, but mutation plays a significant role. Then, from mutation, we may have new alleles arising. The simplest description of such a situation is where we look at the set of *all* alleles that have been present in the population at *some time* for which we have data. A description that encompasses *all* such alleles can then be used from the outset. In this way, we never have to deal with a description that changes dimensionality. Some of the allele frequencies may be zero during some of the time that we have data, but our approach can still be applied. All quantities, such as the force of selection, are well-defined even when some allele frequencies are zero.

Average mutation rates can vary between eukaryotic species ranging from 0.01 × 10^−9^ to 55.58 × 10^−9^ (Wang and Obbard, 2023). However, mutation rates can be substantially higher in certain regions of the genome than in others (Gossmann, Woolfit, and Eyre-Walker, 2011; Nesta, Tafur, and Beck, 2021). We envisage that the primary applications of the present work are to evolutionary trajectories (describing allele or haplotype frequencies) on timescales of the order of tens or hundreds of generations. For such time scales, with typical mutation rates, we expect the influence of mutation on allele frequency trajectories to be very limited. Most new mutations—that are rare—will become lost after a very short time. If, however, mutations are considered to be important, e.g. because the locus of interest lies in a mutational hotspot or the timescale of interest is very long, they can be incorporated into our framework. Recurrent mutations, that transform one allele into another, can be incorporated by adding the term **MX**(*t*) to the right hand side of Eq. (4) (neglecting ‘cross terms’ from selection and mutation, which we assume are very small). New mutations can be incorporated by simply adding more components to the vector of allele frequencies.

Our approach offers estimates of the matrix of fitness-effects, **F**, from allele trajectory data. In this study, we estimated fitness-effects for a genomic region on chromosome 16 of *S. cerevisiae* (and directly from this the matrix of relative fitnesses, **w**). The estimated fitness-effects were quite large in magnitude. In part, this may be attributable to the action of strong selection in a non naturalistic environment during the experimental evolution that was reported. Additionally, the fitness-effects, that we calculated, represent the concatenation of fitness-effects over 30 mitotic generations. We note that two homozygote genotypes (*A*_2_*A*_2_ and *A*_4_*A*_4_) had estimated fitness-effects that correspond to almost complete lethality. However, when estimated values are close to the lowest possible value of the *F*_*i,j*_ they may be sensitive to small numerical errors. We therefore suggest caution in interpreting such results as being effectively lethal, but rather as being strongly deleterious, and susceptible to error. Generally, we expect the fitness-effects of multicellular organisms in natural environments, to which they are reasonably adapted, to be much smaller. Beyond this issue, estimating **F** for four haplotypes allowed us to find evidence for heterozygote advantage, ‘on average’, but also to appreciate variability of the heterozygote fitnesses (some were advantageous, others disadvantageous) that would simply not be resolved within the framework of a biallelic model. With our approach, researchers can address key evolutionary questions about the prevalence of heterosis and the strength of selection that is acting on multiple alleles.

In our analysis of the yeast experimental evolution data, haplotypes were defined within non-overlapping genomic windows of 10 KB (Burke, Liti, and Long, 2014; Phillips et al., 2020). This choice follows the analysis strategy used in the original studies of this experiment (except for some additional thinning), where genomic responses to selection were found to localize to relatively small regions of the genome, typically on the order of 10–20 KB (Burke, Liti, and Long, 2014). In addition, founder haplotype frequencies in these populations were estimated using SNP information aggregated within windows of approximately this size, allowing reliable inference of which founder haplotype occupies a given genomic segment (Phillips et al., 2020). The four founder strains contribute long chromosomal segments to the evolving populations, and recombination during sexual cycles gradually breaks these segments into smaller haplotype blocks. Consequently, a window should not be interpreted as an independent locus under selection, but rather as a local marker of founder ancestry. Selection acting anywhere within a linked genomic region can therefore influence the frequency trajectories of nearby windows through linkage. Because sexual reproduction in the experiment occurred only once every 30 generations, substantial linkage between neighbouring sites is expected to persist over much of the experiment. The inferred fitness-effects should therefore be interpreted as effective fitness differences associated with founder haplotypes within a genomic region, rather than the effects of individual mutations. While a more detailed analysis could explicitly model recombination and reconstruct selected haplotype blocks (Franssen, Barton, and Schlötterer, 2017), the window-based approach provides a practical way to apply the multiallelic framework to genomic time-series data.

As demonstrated, our method can also be applied to genome-wide scans, where **F** is estimated for sliding windows across the entire genome. Using data from an experimental evolution study of *S. cerevisiae* (Burke, Liti, and Long, 2014; Phillips et al., 2020), we observed a widespread pattern in the fitted **F** matrices consistent with heterozygote advantage throughout the genome, which a null-model comparison indicates is unlikely to be explained by drift alone (see Supplementary Material Section 4). Notably, instances where a *homozygote* exhibited the highest fitness appeared to be confined to one highly localized genomic region towards the terminal region of chromosome C16 (see Figure 5). The observed heterozygote advantage may stem from strong artificial selection and/or inbreeding experienced by the original strains prior to the experimental evolution. Consequently, these strains may have undergone intense selection against homozygotes when exposed to a novel environment during the experiment. Interestingly, heterozygote advantage has been reported in other studies on baker’s yeast (Sellis et al., 2016; Aggeli et al., 2022).

In our simulation analyses, for *n* = 4 alleles, trajectories containing at least five sampled time points generally supported stable estimation of the fitness-effect matrix, whereas longer trajectories improved the accuracy of estimated relative fitnesses (see Figure S2). Much of the improvement occurred within the first approximately ten time points, with close convergence in the simulated settings after around 30 time points. Since the number of independent parameters in **F** grows as *n*(*n* + 1)*/*2 − 1, we would expect more time points to be needed for comparably stable and accurate estimation as the number of alleles increases; a systematic characterisation of this scaling is a useful direction for future work. These results provide useful guidance for designing experimental-evolution studies and for selecting suitable empirical time-series datasets, particularly for loci with a similar number of alleles to those studied here.

For instance, our method seems to be suitable for experimental evolution studies involving organisms with short generation times, such as arthropods or nematodes. Importantly, it enables researchers to address diverse questions in evolutionary biology within the typical time frame of a PhD project.

As far as population size, *N* , is concerned, we recommend application of our method to populations of estimated size ≥ 10^4^. This recommendation is supported by our simulation results (Figure S2), which show that the estimation procedure remains accurate at *N* = 10^4^: the mean absolute distance between true and estimated relative fitness values remains below 0.03, and the inferred rank ordering of genotype fitness values closely matches the true ordering after approximately 30 generations. This robustness is likely aided by the use of the Shahshahani metric, which corresponds to a large-*N* approximation of the multinomial sampling likelihood and therefore incorporates stochastic sampling effects to leading order, rather than treating allele-frequency trajectories as purely deterministic. For substantially smaller populations, however, stochastic fluctuations increasingly deviate from this large-*N* approximation, and genetic drift is therefore expected to reduce the accuracy of the inferred fitness matrix.

In practice, this means that the method is particularly well suited to experimental evolution systems in which large population sizes can readily be maintained, such as bacteria and unicellular eukaryotes, including the *S. cerevisiae* populations analysed here. The same considerations apply to genomic time-series data from natural populations. Studies using nucleotide diversity and mutation rate estimates to infer effective population sizes indicate that many natural populations of eukaryotes have effective population sizes ≥ 10^4^ (Gossmann, Keightley, and Eyre-Walker, 2012; Lewin and Eyre-Walker, 2026), suggesting that our method may have broad applicability across natural populations meeting this criterion. Nevertheless, the proposed population-size threshold has thus far been validated only through simulation, and application to empirical time-series datasets from natural populations will require further validation.

Our model is highly relevant for analysing genomic time-series datasets from wild populations, provided sufficiently deep and repeated temporal sampling is available, e.g., as illustrated by our extraction of multiallelic sites from the wild *Drosophila* population data of Kapun et al. (2021), as well as for evolve-and-resequence experiments (Schlötterer et al., 2015). As genomic technologies continue to advance, we anticipate the availability of an increasing number of such datasets in the near future. Notably, our framework could be applied to novel cases, such as where allele frequencies are inferred from sequenced fossils or lake sediments (Decaestecker et al., 2007; Burger et al., 2007) or applied to the analysis of the ongoing evolution of insecticide resistance, e.g. in *Anopheles* mosquitoes (Edi et al., 2014). Furthermore, our model can facilitate genome scans to identify loci under balancing selection. For instance, it can identify loci where the estimated F-matrices make the long-term maintenance of multiple alleles more likely (Lewontin, Ginzburg, and Tuljapurkar, 1978; Siljestam and Rueffler, 2024). This in turn might enable predictions about evolutionary trajectories beyond the observed time frame (Wortel et al., 2023).

There are a number of practical considerations that we wish to highlight when obtaining data on genetic diversity at loci with many alleles.

First, obtaining low frequency variants from population genetic data, by pooling data, requires deep sampling to obtain alleles that are segregating at low frequencies. For example, even at 500X coverage the probability of observing a variant that segregates with a frequency of 0.1 percent (e.g. a new mutant in a diploid population of 500 individuals) is less than 50% (see Figure S8a). Thus, dependent on the expected population size, it might be necessary to sequence a pooled sample at an enormous coverage. However, since sequencing costs have dropped substantially in recent years, deep sequencing may now allow testing of more complex scenarios of selection, due to the identification of rare genetic variants at multiallelic sites.

Second, incorporating low frequency variants is also crucial when considering time-series experiments since this would allow a much improved resolution over time (i.e., avoiding the omission of allele frequency trajectories because they appear hidden at certain time points). This is potentially relevant for evolve and re-sequencing experiments (Schlötterer et al., 2015) or when tracing strains in metagenomic samples over time (Faust et al., 2015; Hildebrand, Moitinho-Silva, et al., 2019; Hildebrand, Gossmann, et al., 2021). However, if low-frequency variation is omitted, e.g., as a consequence of a threshold (Gossmann and Waxman, 2022), then the allele-frequencies of the remaining (above threshold) alleles might be misleading (see Figure S8b). In time-series experiments this may lead to ‘jumps’ that can be misidentified as signatures of rapid change (e.g. selection). In general, as observation thresholds are to be expected due to sampling effects alone it is evident that much diversity and evolutionary dynamics may be missed because it occurs below threshold.

Third, if SNPs are considered in a population-wide sample, then in principle a single base position can, at most, have four alleles. However, we would expect selection to act at the level of DNA sequences that segregate together, and are rarely broken up by recombination events (Williams, 1966), e.g., at the level of haplotypes (Clark, 2004; Fariello et al., 2013), or whole genes (Colbran, Ramos-Almodovar, and Mathieson, 2023). Such natural assemblages will lead, effectively, to a higher number of alleles. Furthermore, when studying epigenetic variation, we can expect an even higher diversity of epialleles because there are more combinations of epigenetic modifications possible than there are unmodified nucleobases (Liu and Zhong, 2024; Kosel et al., 2024). Considering population genetic models with an arbitrary number of alleles not only leads to a realistic description of the variation present, but may also be of high importance, since it may allow capture of all potential targets of selection (Overall and Waxman, 2024). It is possible to obtain haplotype information from short read sequencing data (Garrison and Marth, 2012) and selected haplotype blocks in evolve-and-resequence studies can also be reconstructed from short-read Pool-Seq time-series data by exploiting correlated allele-frequency trajectories (Franssen, Barton, and Schlötterer, 2017). However, a better resolution of haplotypes and potential epimarks can be reached when third generation sequencing technologies are applied, such as PacBio (Rhoads and Au, 2015) or Oxford Nanopore (Wang, Zhao, et al., 2021). A general approach to obtain complex haplotypes might be to use barcoding approaches from long range sequencing data (Sedlazeck et al., 2018).

Fourth, low frequency variation from a population genetic sample might be more difficult to obtain when *individuals* are sampled, which is not uncommon for vertebrate or other multicellular species. Individual based sampling depth will, potentially, need to be moderate in order to identify heterozygous sites, but it would then need to be scaled with the number of individuals tested (see Figure S8a). However, often bioinformatic pipelines take population-wide signatures of polymorphic sites into account, in order to filter out supposed sequencing errors (McKenna et al., 2010; Korneliussen, Albrechtsen, and Nielsen, 2014), which potentially leads to the wrong classification of rare or previously unknown variation. On the other hand, somatic mutations can be mislabeled as germline mutations (Dou et al., 2018; Lupski, 2013; Oota, 2020). Because somatic variation does not propagate to subsequent generations, time-series experiments may be used to identify such types of genetic variation, and therefore may help to accurately identify germline mutations segregating at low frequencies.

## Declarations

### Ethics approval and consent to participate

Not applicable.

### Consent for publication

Not applicable.

### Availability of data and materials

All scripts are available in the github repository https://github.com/NikolasVellnow/selection_multiallelic.

### Author’s contributions

All three authors contributed to the conception of this work. DW carried out the mathematical analysis in consultation with NV and TIG. NV conducted the simulations, in consultation with TIG and DW. All three authors drafted and finalized the manuscript.

## Acknowledgments

Not applicable.

## APPENDICES A Evolutionarily equivalent fitness-effects

In this appendix we relate a property of fitness-effects to a property of relative fitnesses. Relative fitnesses are defined only up to a multiplicative constant. In other words, if *w*_*i,j*_ represents fitness, or relative fitness, of the *A*_*i*_*A*_*j*_ genotype, then with *k* a positive constant, replacing *w*_*i,j*_ by *k* × *w*_*i,j*_, for all *i* and *j*, will produce identical allele frequency trajectories. The corresponding property of the fitness-effects (the *F*_*i,j*_) is that replacing *F*_*i,j*_ by (*k* − 1) + *kF*_*i,j*_, for all *i* and *j*, leads to identical allele frequency trajectories. Thus the matrix of fitness-effects with elements *F*_*i,j*_ is *evolutionarily equivalent* to the corresponding matrix with elements (*k* − 1) + *kF*_*i,j*_.

To show the above, we note the feature that allele frequency dynamics depends only on *ratios* of fitnesses. This allows the use of *relative fitnesses*, which are defined only up to a multiplicative constant. This means if the fitness of the *A*_*i*_*A*_*j*_ genotype is *w*_*i,j*_, then, for all *i* and *j*, replacing *w*_*i,j*_ by *k* × *w*_*i,j*_ where *k* is a positive constant, leads to identical allele frequency dynamics. It also means if, e.g., we designate *A*_1_*A*_1_ as the reference genotype, then we can choose *k* = 1*/w*_1,1_ and the resulting fitness of all genotypes are relative to the fitness of the *A*_1_*A*_1_ genotype.

In terms of the *F*_*i,j*_, the fitness of the *A*_*i*_*A*_*j*_ genotype is proportional to 1+*F*_*i,j*_. An *evolutionarily equivalent* alternative set of fitness-effects, written **F**^*′*^ can be defined, for all *i* and *j*, by

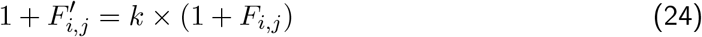

where *k* a positive constant. This equation can be rewritten as

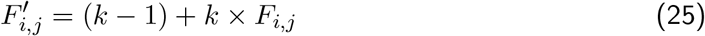

and the 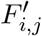 can be used in place of the *F*_*i,j*_ with no effect on allele frequency dynamics.

Additionally, if we, e.g., designate *A*_1_*A*_1_ as the reference genotype, and choose it to have a relative fitness of unity, then this can be taken to correspond to a *vanishing* fitness-effect of this genotype. Given a set of fitness effects, *F*_*i,j*_, where *F*_1,1_≠ 0, using the transformation in Eq. (25) with *k* = 1*/*(1 + *F*_1,1_) leads to a new and evolutionarily equivalent set of fitness-effects 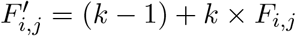 which have the property 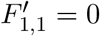.

Such a transformation, which makes a particular genotype the reference genotype, gives it a vanishing fitness-effect. Any *non-lethal genotype* (i.e., that has a positive fitness, or equivalently has a fitness-effect *>* −1) can be converted to the reference genotype.

## B Dynamics of the model

In this appendix we derive the dynamical equation that the frequencies of different alleles obey under a Wright-Fisher model.

We start with an effectively infinite population.

With **X**(*t*) an *n* component column vector containing the frequencies of all *n* alleles in generation *t*, the processes of reproduction and selection of the model described in the “Model” section of the main text lead to the frequency obeying the equation

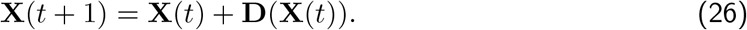

Here **D**(**X**(*t*)) is an *n* component column vector. It depends on the set of frequencies, **X**(*t*), of generation *t* and contains the changes of the frequencies that occur, due to natural selection in generation *t*. In Appendix B we give details of the form of **D**(**x**), which we interpret as the *selective evolutionary force* that acts when the set of allele frequencies is **x**.

In a finite population, apart from the occurrence of reproduction and selection, the process of thinning of the population occurs. This amounts to *non selective sampling* that leads to *N* adults being present at the start of each generation, where *N* is the *census* size. Under the appropriate generalisation of the Wright-Fisher model (Nagylaki, 1992), this has the result that the set of allele frequencies in generation *t* + 1, namely **X**(*t* + 1), does not, generally, coincide with **X**(*t*) + **D**(**X**(*t*)). Rather, **X**(*t* + 1) is distributed *around* the value **X**(*t*) + **D**(**X**(*t*)) according to

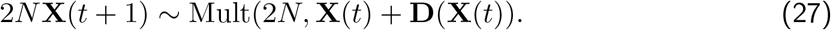

Here Mult(*m*, **p**) denotes a *multinomial distribution*, where the parameter *m* is the number of independent trials, while the parameter **p** is a column vector containing the probabilities of falling into different categories, on each trial^9^.

Let **R**(*t*) denote a multinomial random variable with distribution Mult(2*N*, **X**(*t*) + **D**(**X**(*t*)). The form of **R**(*t*) is an *n* component column vector that has a conditional expectation of

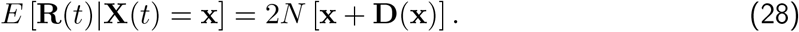

It follows that equivalent to Eq. (27) is the equation **X**(*t* + 1) = **R**(*t*)*/*(2*N*) which we write as

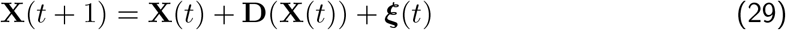

where

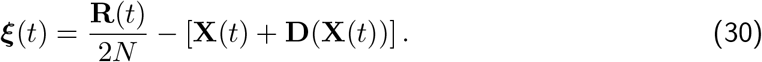

The quantity ***ξ***(*t*) represents statistical fluctuations, due to non selective sampling, around the ‘deterministic prediction’ of **X**(*t* + 1), namely **X**(*t*) + **D**(**X**(*t*)), and ***ξ***(*t*) can be interpreted as a stochastic ‘force’ on allele frequencies that arises from random genetic drift.

From properties of the multinomial distribution (Evans, Hastings, and Peacock, 1993), it follows that the conditional expectation of ***ξ***(*t*) vanishes, i.e.,

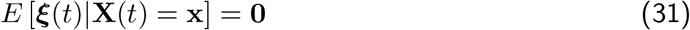

(the right hand side is an *n* component column vector of zeros). Additionally, the variance-covariance matrix of ***ξ***(*t*) is

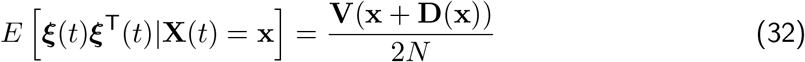

where **V**(**x**) is the matrix

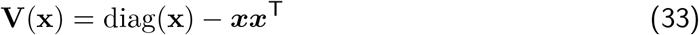

whose elements are given by *V*_*i,j*_(**x**) = *x*_*i*_*δ*_*i,j*_ − *x*_*i*_*x*_*j*_.

On the assumption that all elements of **D**(**x**) are small compared with the corresponding element of **x**, which amounts to saying selection causes a very small *fractional* change in allele frequencies, as commonly holds, we have the standard approximation (Ewens, 2004)

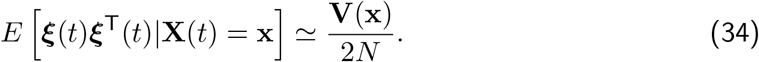

We thus arrive at a description of the dynamics given by Eq. (29) with the primary properties of ***ξ***(*t*) given by Eqs. (31) and (34).

We note that under a diffusion approximation, both time and allele frequencies are approximated as continuous quantities (see e.g., (Kimura, 1955)). This simply entails Eq. (29) becoming *d***X**(*t*) = **D**(**X**(*t*))*dt* + ***ξ***(*t*). The form of ***ξ***(*t*), for continuous time and continuous frequencies, has to have properties that lead to *E* [***ξ***(*t*)|**X**(*t*) = **x**] = **0** and 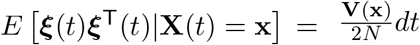. These are closely analogous Eqs. (31) and (34) and can both be achieved by the replacement 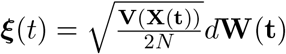 where:

i. **W**(*t*) is a column vector with *n* elements that are ‘Gaussian noise’, i.e., independent Wiener processes (Tuckwell, 2019);
ii. the square root of the matrix **V**(**x**) in 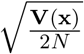 denotes the *principal square root*. Since **V**(**x**) is the variance covariance matrix of a multinomial random variable it is real, symmetric and positive semidefinite, with a smallest eigenvalue of zero, and a bounded largest eigenvalue. The matrix 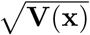 that is the principal square root also has these properties (see e.g., Higham, 2008). The principal square root converts the ‘noise’ *d***W**(**t**) into real changes in allele frequencies, which sum to zero.

Lastly, deviations of a population from being ‘ideal’ are often incorporated into the diffusion approximation by replacing the census population size, *N* , by an ‘effective’ population size *N*_*e*_ (Wright, 1931). This leads to **X**(*t*) obeying the stochastic differential equation 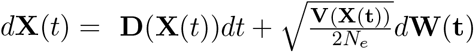.

## C Deriving the form of the selective force

In this appendix we provide an elementary derivation of a compact representation of the evolutionary force associated with selection.

We have *n* alleles labelled *A*_*i*_ with *i* = 1, 2, … , *n* and assume the *A*_*i*_*A*_*j*_ genotype has a fitness proportional to 1 + *F*_*i,j*_ with *F*_*i,j*_ = *F*_*j,i*_. Representing fitness in terms of *F*_*i,j*_ is not standard, hence we derive all results from first principles. Results in terms of fitness written as *W*_*i,j*_ are well known in the literature (Bürger, 2000; Ewens, 2004).

Let *x*_*i*_ be the frequency of allele *A*_*i*_ in adults of a particular generation, and let 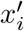 be its frequency in offspring after random pairing, reproduction, and selection.

Contributions to 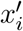 arise from: (i) offspring with the *A*_*i*_*A*_*i*_ genotype and (ii) offspring with the *A*_*i*_*A*_*j*_ genotype, with *j*≠ *i*. With all sums running from 1 to *n* unless otherwise stated, this leads to

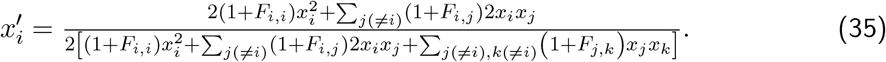

This can be simplified to

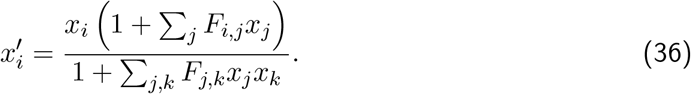

Alternatively, we have

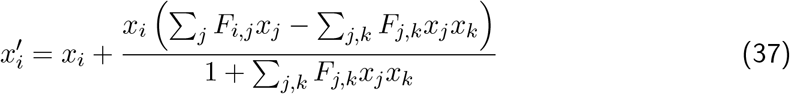

and we rewrite this equation as

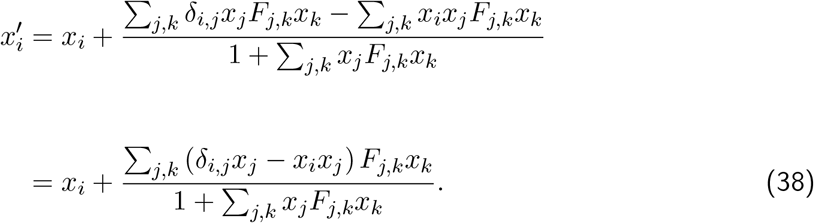

Using [**y**]_*i*_ to denote the *i*’th element of the vector **y**, and with **V**(**x**) the *n* × *n* matrix with elements

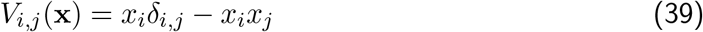

we can write Eq. (38) as 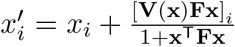 where **F** is a matrix with elements *F*_*i,j*_. From this we obtain

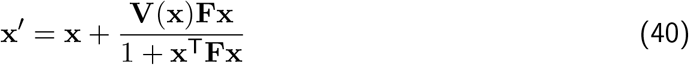

and this equation allows us to identify the second term on the right hand side with the evolutionary force of selection, i.e., we identify

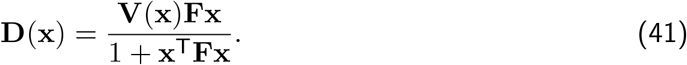

Such a result is implicit in the mathematical work of Akin (1990).

While we could formally represent **D**(**x**) as **V**(**x**) times the gradient of 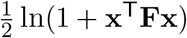 ln(1 + **x**^T^**Fx**), we do not do so since such a representation does not generally hold when **F** is frequency-dependent.

## D Estimating the F matrix

Let the exact fitness of the *A*_*i*_*A*_*j*_ genotype, for all *i* and *j*, be proportional to 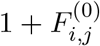 and let **F**^(0)^ denote the *n* × *n* symmetric matrix containing the 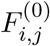. In this appendix we give a method for estimating **F**^(0)^ from allele frequency trajectory data via an optimisation procedure.

We present the method under the assumption that **F**^(0)^ is independent of allele frequencies. Extensions, to e.g., frequency dependent selection, may be possible if an **F**^(0)^ matrix with a specific frequency-dependent form is assumed/adopted, and parameters within this form are included within the optimisation procedure.

We shall first present a method of estimating **F**^(0)^ when allele frequencies are known at consecutive generations. We then extend this to the case where allele frequencies are only known for particular, non-consecutive, generations.

### D.1 Allele frequencies known every generation

We begin with data for a locus with *n* alleles. The data takes the form of a trajectory

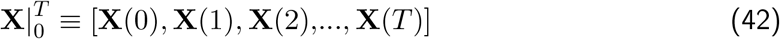

where for *t* = 0, 1, …, *T* , the quantity **X**(*t*) is a column vector of length *n*, and contains *n* allele frequencies. Thus 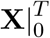 is an *n* × (*T* + 1) matrix.

Under purely selective dynamics, as described in the main text (Eq. (26)), the behaviour of **X**(*t*)) is given by^10^

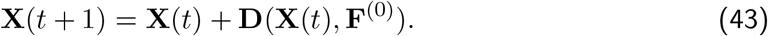

The form of the matrix **F**^(0)^ is unknown and our objective is to estimate it from the *data* (i.e., from the trajectory, 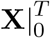).

To proceed, we define another trajectory

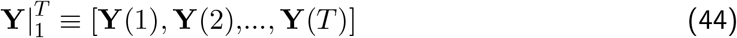

which is a matrix of size *n* × *T* . We determine 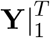 from the known trajectory data (i.e., 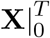) and an arbitrarily chosen **F** matrix via

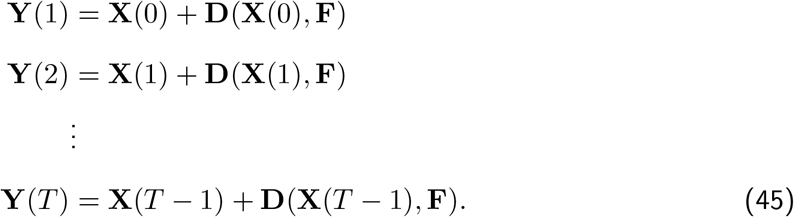

That is, each **Y**(*t*) is determined from: (i) the data frequencies of one generation earlier, namely **X**(*t* − 1), and (ii) the **F** matrix. We can say **Y**(*t*) is the *predicted value* of **X**(*t*), given **X**(*t* − 1) and the matrix **F**.

We now define a ‘cost function’ *C*(**F**) that is minimised when **F** = **F**^(0)^.

Let **X**(*t*) denote a column vector of observed allele frequencies at time *t*, with elements *X*_*i*_(*t*) (*i* = 1, 2, …, *n*).

Let **Y**(*t*) denote a column vector of predicted allele frequencies at time *t*, with elements *Y*_*i*_(*t*) (*i* = 1, 2, …, *n*), that are based on **X**(*t* − 1) and the matrix **F** of fitness-effects.

A basic form of the cost of **F** is

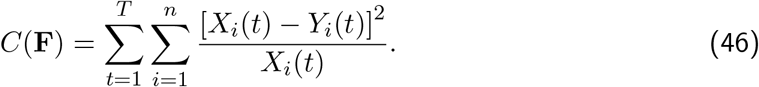

The weightings [*X*_*i*_(*t*)]^−1^ in Eq. (46) correspond to the Shahshahani metric (Shahshahani, 1979; Akin, 1990), and originate in the inverse of the matrix **V**(**X**(*t*)), which, up to a factor proportional to *N*_*e*_ is the natural weighting of allele frequency changes when subject to random genetic drift. Indeed a log likelihood function based on this weighting is proportional to *C*(**F**).

We note that when **Y**(*t*) = **X**(*t*) the cost vanishes:

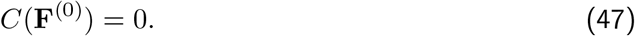

The cost also vanishes for **F** matrices that are *evolutionarily equivalent* to **F**^(0)^ (see Appendix A) and hence produce the same allele frequency trajectories. For any **F** that is not evolutionarily equivalent to **F**^(0)^ we have *C*(**F**) *>* 0 since the **Y**(*t*) will not generally coincide with the **X**(*t*).

When we carry out the minimisation, we require that two points are satisfied.

i. The matrix **F** is unambiguous. We achieve this by always taking *F*_1,1_ = 0, which sets *A*_1_*A*_1_ to be the reference genotype, and the fitnesses of all genotypes are then given relative to the fitness of this genotype (see Appendix A).
ii. When changes of the matrix **F** are made, it is always kept a symmetric matrix. When there are *n* alleles, there are *n*(*n* + 1)*/*2 distinct elements of **F**. On Satisfying point reduces this to *n*(*n* + 1)*/*2 − 1 independent elements. We satisfy point (ii) by expressing results in terms of a vector **v** with *n*(*n* + 1)*/*2 − 1 elements, and using a function **S**(**v**) that produces, from the elements of **v**, a symmetric matrix with a vanishing (1, 1) element. Thus the basic form of the procedure carried out is a minimisation of *C*(**S**(**v**)) over **v**. With 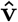 the vector that minimises *C*(**S**(**v**)), the estimate of **F**^(0)^ is 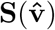.

While the basic form of the cost function in Eq. (46), associated minimisation procedure is theoretically justified, in practise we have found it necessary to make two modifications.

The first modification is to add a numerically small number, *α*, to the allele frequency in the denominator, in order to avoid infinite weightings associated with vanishing allele frequencies.

The second modification is to add a ‘ridge penalty’ (Hoerl and Kennard, 1970) which is an accepted method of suppressing large values in the solution. The ridge penalty biases the elements towards zero and reduces variance. It takes the form of an additional term in the cost. We adopted a term in the form of the squared length of the vector **v**.

Thus the fitness function adopted in this work is

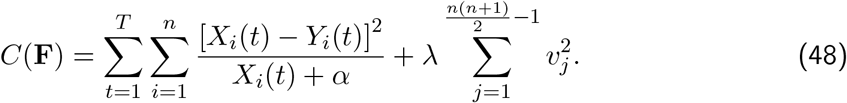

In practise, we found that *α* = 10^−6^ and *λ* = 6 × 10^−3^ gave acceptable results. We assessed the sensitivity of estimation accuracy to these two hyperparameters systematically, across a range of population sizes (see Supplementary Material). We find little sensitivity to *α*, whereas the best-performing *λ* decreases with population size; the values adopted here fall within a region of comparably good performance across all sizes tested.

For empirical trajectories minimisation of the cost function yields the fitness-effect matrix that best reproduces the observed frequency changes, according to the model specified.

### D.2 Allele frequencies not sampled every generation

We now assume the allele frequencies are only known in generations *T*_1_, *T*_2_, …, *T*_*g*_ where generally *T*_*i*+1_ − *T*_*i*_≠ 1 and this interval may be variable. We define the ‘sampled’ trajectory by

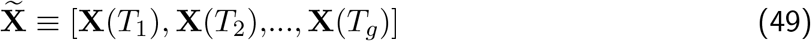

which is a matrix of size *n* × *g*.

To proceed, we define another sampled trajectory

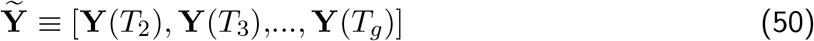

which is a matrix of size *n* × (*g* − 1). We determine 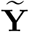 from 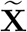 and an arbitrarily chosen **F** matrix, with the calculations as follows.

To determine **Y**(*T*_2_) ≡ **Y**(*T*_1_ + (*T*_2_ − *T*_1_)) we carry out the *T*_2_ − *T*_1_ calculations^11^

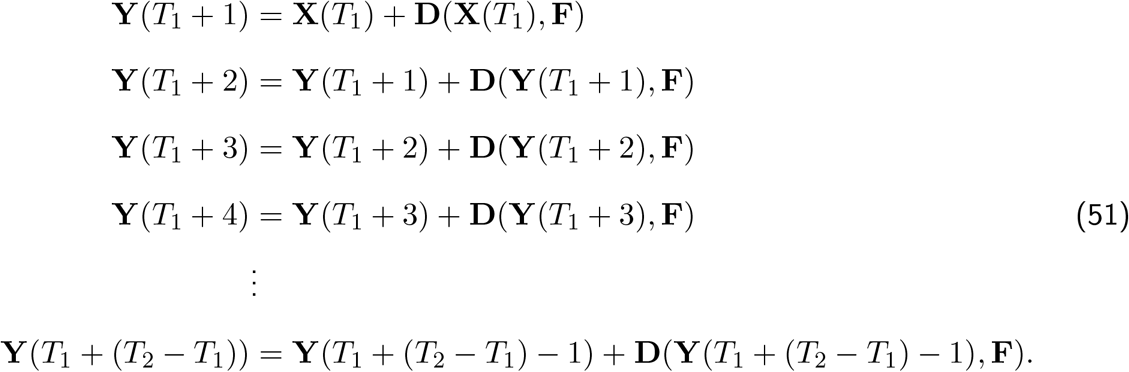

Equivalently, to determine **Y**(*T*_3_) ≡ **Y**(*T*_2_ + (*T*_3_ − *T*_2_)) we carry out the *T*_3_ − *T*_2_ calculations

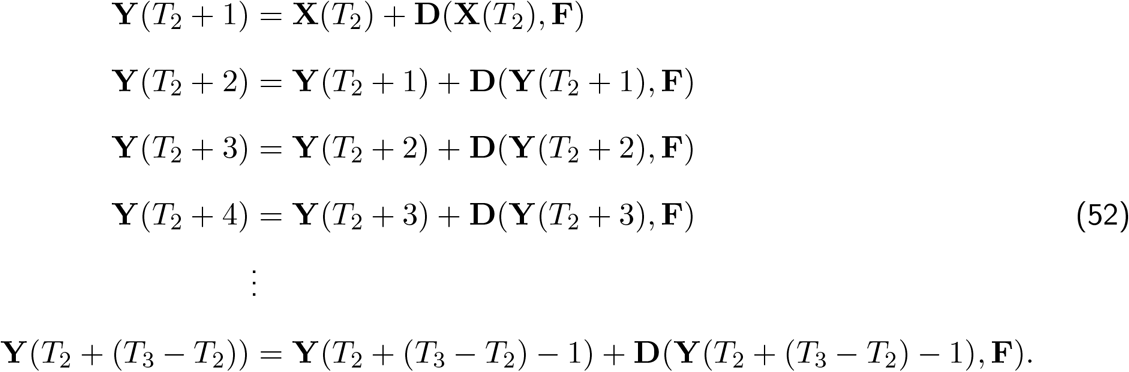

We proceed in this way until we have calculated all **Y**(*T*_1_), **Y**(*T*_2_), …, **Y**(*T*_*g*_).

The ‘cost’ of matrix **F** is then taken to be

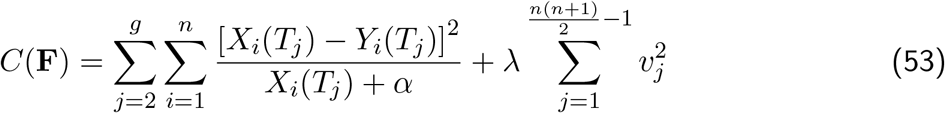

where, as in Section D.1, we parameterised **F** in terms of an *n*(*n* + 1)*/*2 − 1 component vector **v**, and used a function **S**(**v**) to construct **F** from **v**:

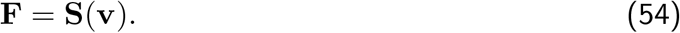

The minimisation that is carried out is that of *C*(**S**(**v**)) over **v**. As before, with 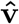 the vector that minimises *C*(**S**(**v**)), the estimate of **F**^(0)^ is 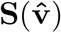.

## Supplementary Material

In Sections 1 and 2 of this Supplementary Material we give details of estimating fitness-effect matrices from empirical frequency trajectories, including an additional example illustrating a case where the assumption of constant selection is poorly supported. In Section 3 we explore how trajectory length, genetic drift, and the ridge-penalty hyperparameters influence the estimate of the fitness-effect matrix. In Section 4 we provide simulation-based validation of our empirical yeast findings, including a comparison against a neutral, drift-only null model and an assessment of the reliability of our heterozygote-advantage classification. In Section 5 we explore the sensitivity of the number of alleles detected at a locus to the level of filtering applied to pooled sequencing data, and in Section 6 we provide calculations and simulations to illustrate detection and threshold effects.

## 1 Details of estimating the F matrix from empirical data

For the estimation of **F** with our ‘cost function’ *C*(**F**) (see Eq. 53 in Appendix D.2), we used the ‘fmincon’ optimisation function in MATLAB R2026a (The Mathworks Inc., 2025), which implements gradient-based methods for constrained minimization. This method performed well, especially in combination with a ridge regularization with a *λ* = 6 × 10^−3^, which penalizes very extreme and thereby unlikely fitness values.

## 2 Estimating F matrix from empirical data: Another yeast example

As another example, we also estimated **F** (with constrained minimization) for trajectories of the haplotypes in the 10 KB window around position 436,424 on chromosome C12, for replicate population 3 (see Figure S1a), from the data set of Burke, Liti, and Long (2014) and Phillips et al. (2020). The estimated **F** and the corresponding matrix of relative fitnesses **w** (minimum cost: 1.276) are:

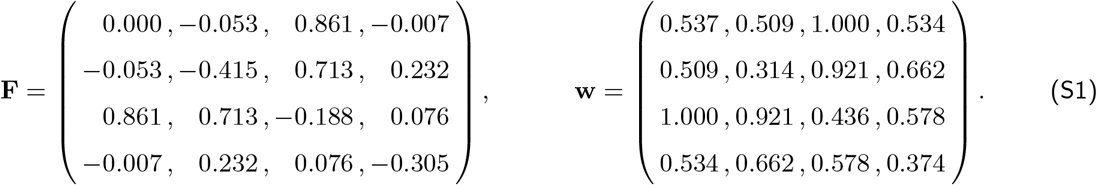

The estimated **F** leads to trajectories that resemble the empirical ones, although the resemblance is not very strong (see Figure S1b). In particular, we hypothesize that after generation 9 the selective forces may have changed; since under that scenario **F** would not have been constant, our estimation method may not find a well-matching **F**, consistent with this window’s comparatively high cost value of 1.276, which lies above the 75th percentile (0.785) of the cost-value distribution across all 1, 161 yeast windows. More broadly, this distribution is tightly concentrated at low values for the majority of windows (median 0.550; interquartile range [0.421, 0.785]), but has a long right tail (mean 1.217; 95th/99th percentiles 4.716/12.870). This indicates that while most windows are well described by a constant **F**, a minority – such as the window around position 436,424 on chromosome C12 – show an elevated cost, plausibly reflecting genuine violations of the constant-selection assumption rather than a general shortcoming of the estimation procedure.

**Figure S1.**
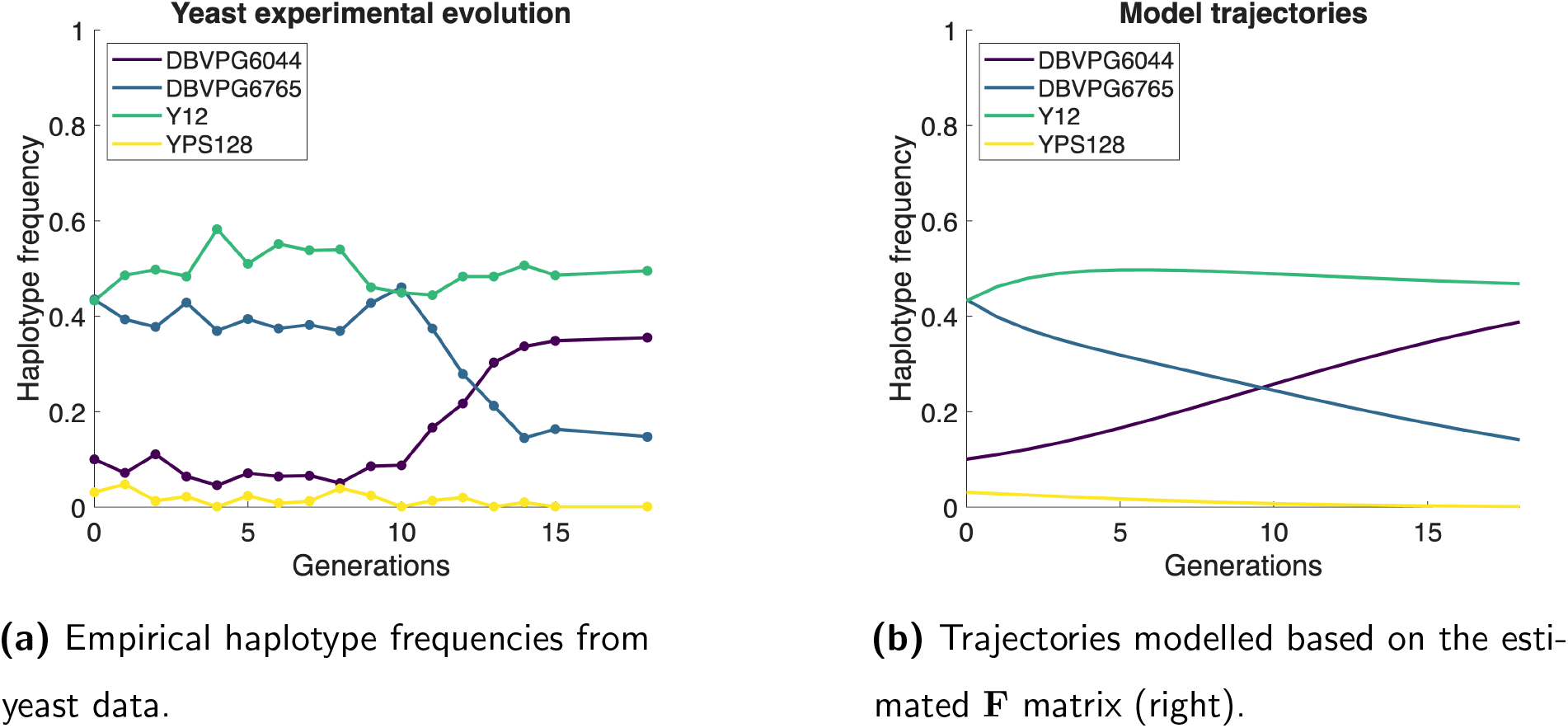
Estimating the F matrix from brewer’s yeast haplotype frequency trajectories. Data is from a 10 KB window around position 436,424 on chromosome C12 for replicate 3 (Burke, Liti, and Long, 2014; Phillips et al., 2020). Haplotypes are DBVPG6044, West African in dark blue, DBVPG6765, wine/European in purple, Y12, sake/Asian in green, and YPS128, North American in yellow.

## 3 Performance of the inference method across population sizes and trajectory lengths

In order to investigate how the length of allele frequency trajectories affects the estimate of **F** and **w** correspondingly, we proceeded as follows.

We generated trajectories based on a known **F**^(**0**)^. Using trajectories produced in this way, but of different length (i.e., different time duration), yielded different estimates of the **F** matrix. We compared the estimates with the actual fitness effect matrix, namely **F**^(**0**)^.

We began by generating an **F**^(**0**)^ with elements, drawn from a normal distribution with mean 0 and standard deviation 0.1, along with random initial allele frequencies. From these, we simulated trajectories with varying population sizes (*N* = ∞, *N* = 10^6^, *N* = 10^5^ and *N* = 10^4^) of length 40 generations. We next used our method to estimate **F** and **w**, based on trajectories of different lengths (corresponding to different experimental observation periods), ranging from 1 to 40 generations. Next, we calculated, for each estimated **w** matrix, the distance to the true relative fitness matrix (namely **w**^(**0**)^), along with the corresponding Spearman rank correlation coefficient.

In Appendix D.1 we introduced the **v** vector, which contains the independent elements of the matrix of fitness effects. The distance was calculated as the mean absolute error (MAE) between entries of the true **w**^(**0**)^ and the estimated **w**:

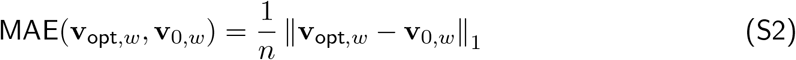

where ∥ · ∥_1_ represents the *ℓ*_1_- norm, and **v**_0,*w*_ and **v**_opt,w_ represent the true and the estimated vector **v**, respectively, when transformed to relative fitness values, and *n* represents their length.

Finally we repeated this procedure for 256 independently generated **F**^(**0**)^ matrices. From these matrices, for each of the different observation lengths, we calculated the median, along with the 25% and 75% quantiles of MAE.

We observe good convergence of the distance and correlation measures within 40 generations (see Figure S2, row 1-2). A substantial part of the improvement in the **w**-estimation occurs within the first 10 generations and a very well estimated **w** is reached after 30 generations (see Figure S2, row 1). The rank ordering of estimated fitness values correlates strongly with the true ordering after about 10 generations and exceeds the null expectation after roughly 30 generations (see Figure S2, second row). This suggests that statements about the rank ordering of estimated fitness values of different genotypes will typically be correct after relatively short observation periods. By contrast, the error of the precise values of **w** are more influenced by stochasticity introduced by genetic drift, but are expected to be smaller than 0.03 even for rather small populations like *N* = 10^4^.

Furthermore, we also simulated trajectories for the same set of population sizes, but under neutral evolution, where all genotypes had the same fitness and used our method to estimate **w**. We then calculated the mean relative fitness for each estimated **w** in replicate trajectories and plotted the change over time and the distribution at the end of 40 generations (Figure S2, row 3 and 4, respectively). If perfectly estimated, the mean relative fitness should be 1.0. As can be seen from Figure S2, row 3 the estimated values lie close to the true value. Comparing the mean relative fitness inferred from real data with the 5th percentile of this null distribution provides a simple statistical test for rejecting the hypothesis of neutral evolution.

**Figure S2.**
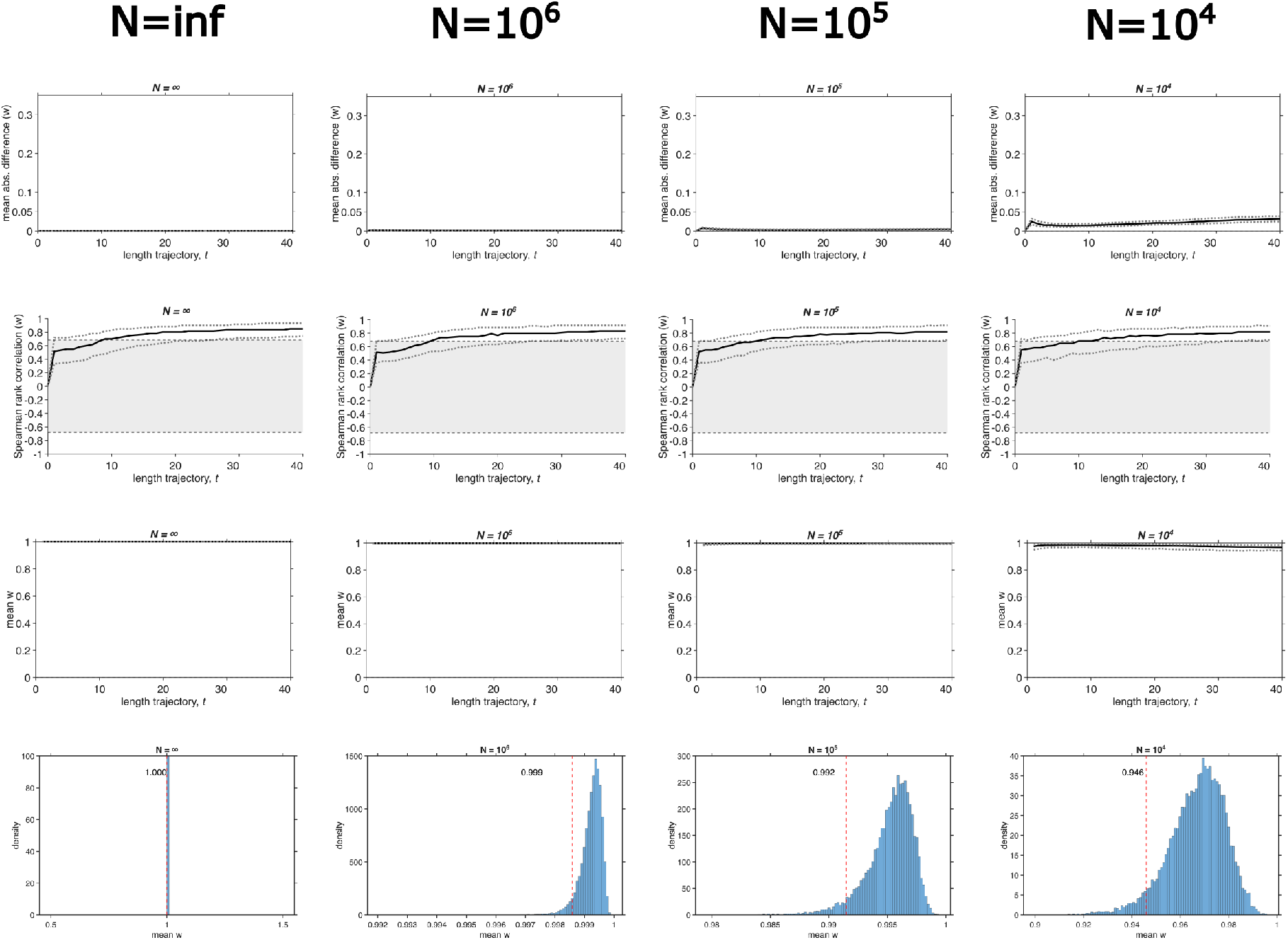
Performance of the inference method across population sizes and trajectory lengths. Columns correspond to population size *N* , and rows show different summary measures or selection scenarios. For the first three rows, the x-axis indicates the length of the observed trajectory. The first row shows the mean absolute difference between the true and the estimated relative fitness matrix. The second row shows the Spearman rank correlation coefficient between the true and estimated **w** matrices; the grey shaded region indicates the 95% confidence interval expected for the correlation between two random matrices (null expectation). The third row shows 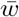, the mean relative fitness per estimated matrix under neutral evolution, representing the method’s null expectation in the absence of selection. In rows 1–3, the black line denotes the median across 256 replicate simulations and the dotted lines indicate the corresponding percentile ranges (25–75% in rows 1–2, 2.5–97.5% in row 3). The bottom row shows the distribution of 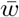 after 40 generations of neutral evolution across 16,384 replicate simulations; the dashed red line marks the 5th percentile, below which 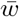 values are unlikely under neutrality.

Furthermore, we also simulated trajectories for the same set of population sizes, but under neutral evolution, where all genotypes had the same fitness, and used our method to estimate **w**. For each replicate trajectory, we then calculated the mean of the entries of the estimated **w**, denoted 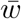, and plotted its change over time and its distribution at the end of 40 generations (Figure S2, row 3 and 4, respectively). If perfectly estimated, 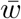 should be 1.0. As can be seen from Figure S2, row 3, the estimated values lie close to the true value. Comparing 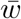 inferred from real data with the 5th percentile of this null distribution provides a simple statistical test for rejecting the hypothesis of neutral evolution.

### Sensitivity of estimation to the ridge-penalty hyperparameters *α* **and** *λ*

The cost function used for estimating **F** (Eq. 48 in Appendix D) includes two hyperparameters: *α*, a small constant added to the denominator to avoid infinite weightings at vanishing allele frequencies, and *λ*, the ridge-penalty weight that shrinks estimated fitness effects towards zero. In the main text we used *α* = 10^−6^ and *λ* = 6 × 10^−3^, values chosen based on empirical performance. Here we assess the sensitivity of estimation accuracy to these choices systematically, and across population size.

For each combination of *α* ∈ {10^−8^, … , 10^−2^} (7 log-spaced values) and *λ* ∈ {10^−8^, … , 10^−1^} (24 log-spaced values), and for each of three population sizes (*N* = 10^3^, 10^4^, 10^6^), we generated *n*_rep_ = 64 independent replicate simulations. In each replicate, we constructed a random 4-allele fitness matrix **F**^(0)^ by drawing its independent entries i.i.d. from a normal distribution (mean 0, SD 0.1), and simulated a population of the given size, starting from random initial allele frequencies, evolving under this true **F**^(0)^ for *T* = 100 generations. For each replicate, we then estimated **F** (and the corresponding relative fitness matrix **w**) from the full simulated trajectory, using the given (*α, λ*) pair, via the same multi-start constrained optimization procedure used elsewhere (*n*_starts_ = 80).

For each replicate we computed two measures of estimation accuracy, applied to **w**: the Spearman rank correlation between the true and estimated relative fitness vectors, and the mean absolute error (MAE) between them (as defined in Eq. S2). We then summarized these measures, at each (*α, λ*) combination and population size, by their mean across the 64 replicates, yielding six 7 × 24 grids of estimation accuracy across the hyperparameter space – one pair of grids (rank correlation, MAE) per population size (Figure S3).

We observe little sensitivity to changes in *α* across the analysed values; performance instead depended primarily on *λ*, and the best-performing *λ* decreased with population size, from *λ* ≈ 5 × 10^−2^ at *N* = 10^3^ to *λ* ≈ 3.7 × 10^−4^ at *N* = 10^6^. The default values used in the main text (*α* = 10^−6^, *λ* = 6 × 10^−3^) fall within a region of comparably good performance at all three population sizes, and can, therefore, be considered a good compromise in situations where the precise population size is unknown.

**Figure S3.**
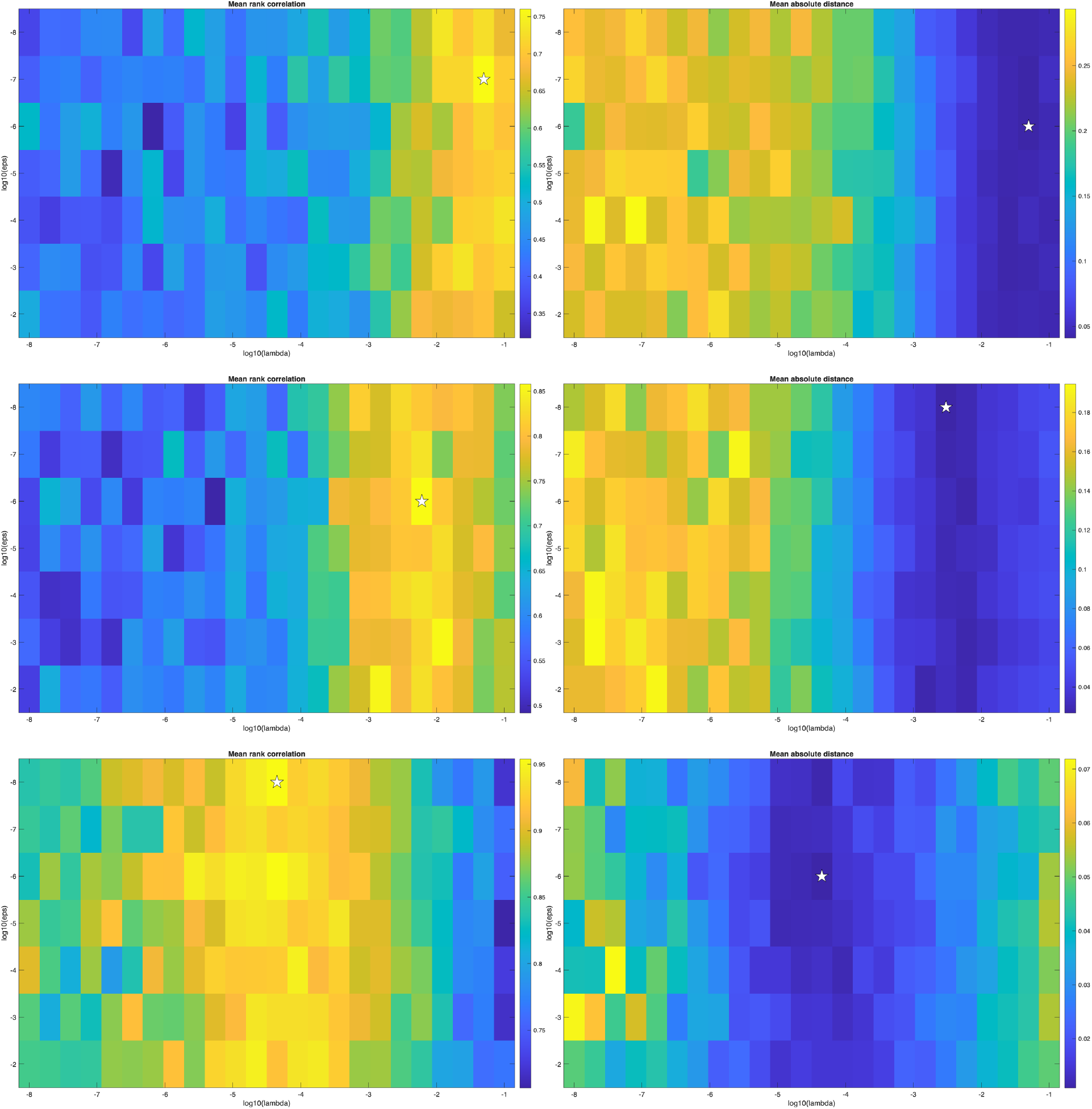
Estimation accuracy is robust across a wide range of the ridge-penalty hyper-parameters *α* and *λ*, across population sizes. Heatmaps show estimation accuracy across a grid of *α* (7 log-spaced values, 10^−8^ to 10^−2^) and *λ* (24 log-spaced values, 10^−8^ to 10^−1^), summarized across *n*_rep_ = 64 replicate simulations per (*α, λ*) combination, at *N* = 10^3^ (top row), *N* = 10^4^ (middle row), and *N* = 10^6^ (bottom row). Left column: mean Spearman rank correlation between the true and estimated relative fitness vectors. Right column: mean absolute error (MAE) between the true and estimated relative fitness vectors. In all heatmaps, colour indicates the value of the corresponding metric, from dark blue (low) to yellow/orange (high); since high rank correlation and low MAE both indicate good estimation accuracy, yellow/orange denotes good performance in the left column, while dark blue denotes good performance in the right column. In each panel, the white star marks the (*α, λ*) combination that performed best by the corresponding metric.

## 4 Validation of yeast findings with simulated data

### Non-neutral evolution

To test whether neutral evolution due to genetic drift could cause relative fitness values as high or higher than we have found for the yeast data, we simulated evolutionary trajectories for relatively low population sizes (*N* = 10^3^, 10^4^) and compared the results with the empirical yeast data. We set the true 4×4 fitness matrix **F**^(0)^ to be exactly neutral (all entries zero), so that no true fitness differences existed among the genotypes. For each of *n*_rep_ = 2^14^ = 16,384 independent replicate simulations at each population size, we drew random initial allele frequencies and simulated genotype frequencies forward for *T*_max_ = 40 generations under Hardy–Weinberg random mating and viability selection governed by **F**^(0)^. Since **F**^(0)^ = **0**, selection left genotype frequencies unchanged at every generation, so any change in allele frequencies arose purely from genetic drift implemented via multinomial sampling of *N* diploid individuals’ genotypes each generation rather than from true fitness differences. We then applied the same estimation procedure used for the empirical yeast data and the random-**F**^(0)^ simulations described below (multi-start constrained optimization of the ridge-penalized cost function; *n*_starts_ = 80, *λ* = 0.0061) to each trajectory, yielding an estimated fitness matrix **F**, which we converted to the relative fitness matrix **w** (as defined above) and summarized via its mean 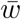: values close to 1 indicate near-uniform estimated fitness, as expected under true neutrality, while low values indicate spuriously inferred fitness differences. Repeating this across all replicates yielded, for each population size, a null distribution of 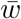 attributable entirely to genetic drift and estimation noise, against which we compared the corresponding empirical distribution of 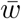 across the yeast genomic sites (computed identically from each site’s estimated **F**).

At *N* = 10^4^, the null distribution had a median 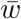 of 0.968 (5% quantile: 0.946); at *N* = 10^3^, drift alone pushed the null median down to 0.865 (5% quantile: 0.796), reflecting the expected increase in drift at smaller population size. The empirical median 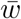 across yeast sites was 0.644 – substantially lower than either null distribution, including the more permissive *N* = 10^3^ case (Figure S4). Correspondingly, the large majority of empirical sites fell below the lower 5% quantile of the null distribution at both population sizes: 99.57% (1156 of 1161 sites) at *N* = 10^4^, and 98.54% (1144 of 1161 sites) at *N* = 10^3^.

This indicates that the pronounced fitness heterogeneity observed among yeast genotypes is highly unlikely to be explained by genetic drift alone. Note that our cost function already incorporates a large-*N* approximation to the sampling likelihood under drift, via the Shahshahani metric, and the population sizes used here (*N* = 10^3^, 10^4^) are well below the yeast experiment’s reported effective population size (*N*_*e*_ ≈ 10^6^; see the more detailed discussion of this quantity in the following subsection), providing a conservative test of this approximation under stronger drift than is likely present in the real data.

**Figure S4.**
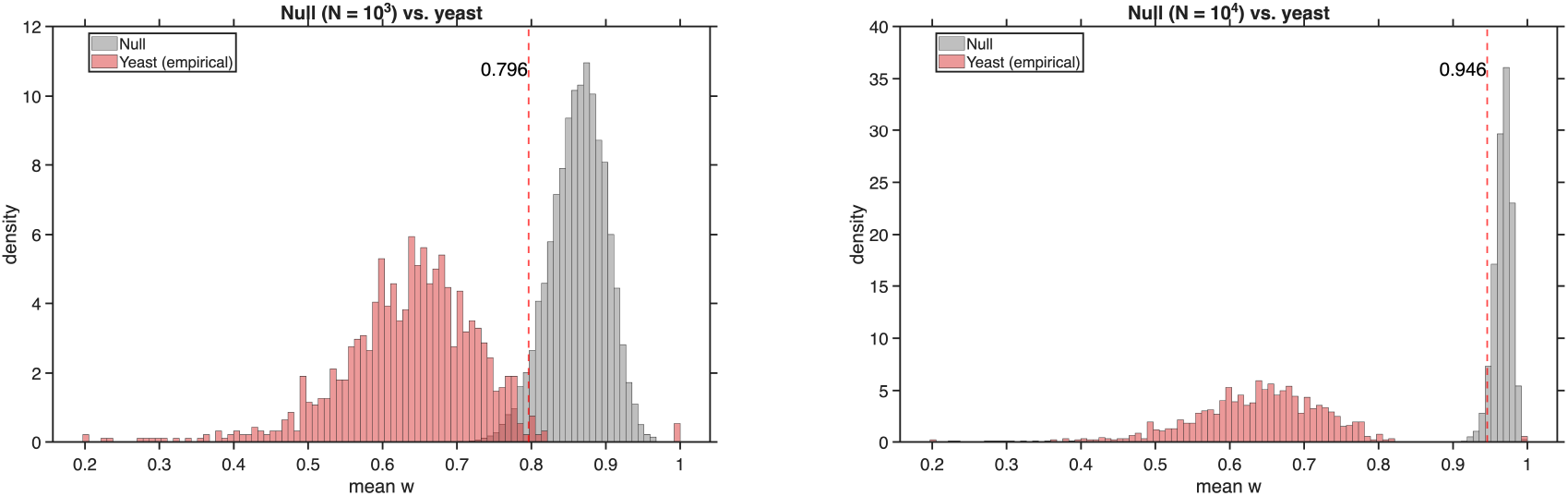
Empirical yeast fitness heterogeneity exceeds the neutral null expectation at both population sizes. Distributions of the mean relative fitness 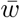 under simulated neutral evolution (grey; *n* = 16,384 replicates per panel) compared to the empirical distribution of 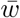 across the 1161 yeast genomic sites (red), at *N* = 10^3^ (left) and *N* = 10^4^ (right). The dashed red line marks the 5th percentile of the null distribution in each panel. The empirical yeast distribution lies almost entirely below this threshold at both population sizes (98.54% and 99.57% of sites at *N* = 10^3^ and *N* = 10^4^, respectively), indicating that the fitness heterogeneity observed in yeast is highly unlikely to arise from genetic drift alone.

### Heterozygote advantage

To test whether our estimation method could itself produce a signal of heterozygote advantage that is not actually present in the underlying fitness matrix—and how reliably it detects heterozygote advantage when it is present—we simulated evolutionary trajectories in MATLAB and applied our estimation procedure to the simulated data.

For each of three population sizes (*N* = 10^3^, 10^4^, 10^5^), we independently generated 2^14^ = 16,384 replicate simulations (a number chosen for computational convenience). In each replicate, we constructed a random 4-allele fitness matrix **F**^(**0**)^ by drawing its entries from a normal distribution (mean 0, SD 0.1); one diagonal entry is fixed at 0 as a reference allele. We then simulated a population with four alleles, starting from random initial frequencies, evolving under this true **F**^(**0**)^ for 40 generations, and applied our estimation procedure to the full simulated frequency trajectory to obtain an estimated **F**.

To assess the reliability of this result, we repeatedly subsampled from the pool of 16,384 replicates. For each population size, we drew 1000 independent samples, each consisting of 1161 replicates chosen at random without replacement (i.e., no replicate is counted twice within a given sample). Because each of the 1000 samples is drawn afresh from the full pool of 16,384, the same replicate can appear in multiple samples. Sample size (1161) was chosen to match the number of analysed genomic windows in the yeast experiment. Within each sample, we split the replicates into two groups based on the <u>true, simulated</u> **F**^(**0**)^: replicates where the most-fit genotype was a homozygote, and replicates where it was a heterozygote. We then computed, within each group, the proportion of replicates where the <u>estimated</u> **F** also indicated a heterozygote as most-fit. This yields, for each population size, both a false-positive rate (spurious heterozygote calls when the truth is homozygous) and a true-positive rate (correct heterozygote calls when the truth is heterozygous) for our estimation procedure.

At every population size tested, our procedure showed substantial discriminative power between true heterozygote advantage and its absence: a heterozygote call was 1.9–3.9 times more likely when the true most-fit genotype was genuinely a heterozygote than when it was a homozygote (positive likelihood ratio TPR*/*FPR: 1.86, 3.28, and 3.87 at *N* = 10^3^, 10^4^, and 10^5^, respectively). This discriminative power increased with population size, driven primarily by a false-positive rate that fell sharply with *N* – from 0.508 (95% CI [0.459, 0.556]) at *N* = 10^3^ to 0.278 (95% CI [0.233, 0.323]) at *N* = 10^4^ and 0.232 (95% CI [0.191, 0.276]) at *N* = 10^5^ – while the true-positive rate remained consistently high throughout, at 0.945 (95% CI [0.930, 0.960]), 0.912 (95% CI [0.892, 0.932]), and 0.897 (95% CI [0.876, 0.918]), respectively (Figure S5). The effective population size of *N*_*e*_ ≈ 10^6^ reported for this yeast experiment reflects a bottleneck size deliberately imposed at two separate points within each 30-generation cycle between forced outcrossing events – once several generations after mating, and again mid-way through the subsequent liquid-culture growth phase – with substantial population expansion (to stationary-phase densities of 0.8–4 × 10^9^ cells/ml) occurring between bottlenecks (cf. Fig. S1 of Burke, Liti, and Long, 2014). Consequently, *N*_*e*_ ≈ 10^6^ does not correspond directly to the population size at a single, uniform generation in our simulations, which model one full cycle as a single discrete generation. Instead, the effective population size relevant to genetic drift over such a cycle depends on the bottleneck size, the number of bottlenecks per cycle, and the extent of intervening growth, and cannot be recovered from the bottleneck size alone. We therefore do not attempt to map *N*_*e*_ ≈ 10^6^ onto a specific point in our simulated range, noting only that our tested population sizes (*N* = 10^3^–10^5^) are comparable to or smaller than the bottleneck size itself, giving some indication of expected performance under repeated strong bottlenecks. This makes it unlikely that the high prevalence of heterozygote advantage observed in the yeast data is primarily driven by estimation artifacts.

**Figure S5.**
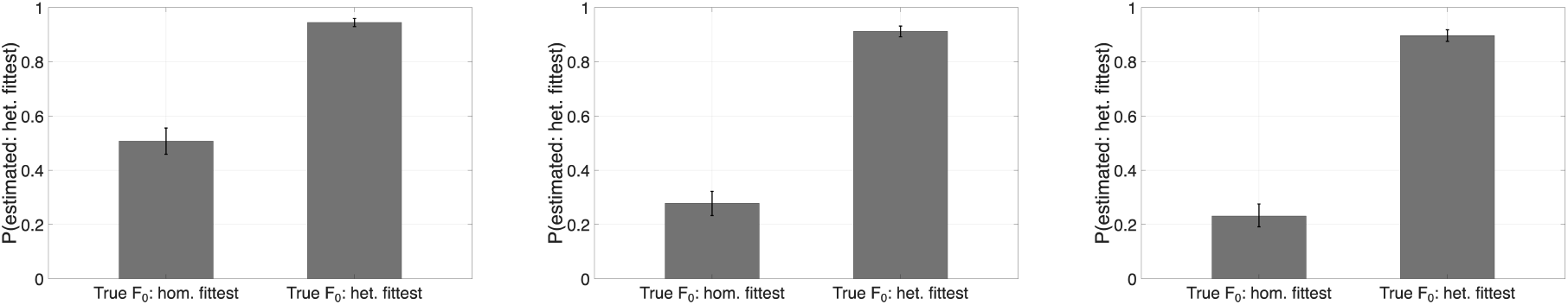
The estimation procedure discriminates heterozygote advantage from its absence, though with a non-negligible false-positive rate at small population size. For each population size (*N* = 10^3^, 10^4^, 10^5^; left to right), replicates were split by the true identity of the most-fit genotype, and we computed the proportion of replicates for which the estimated **F** also called it a heterozygote. The left bar is thus a false-positive rate; the right bar a true-positive rate. The false-positive rate decreases with population size (0.508 → 0.278 → 0.232) while the true-positive rate remains consistently high (0.945 → 0.912 → 0.897). Error bars denote 95% CI over subsamples (see main text).

Both analyses in this section rely on subsampling procedures that treat the 1,161 replicates within each simulated sample as statistically independent. The corresponding empirical genomic windows, however, are not independent: as discussed in the main text, adjacent 10 KB windows may retain substantial linkage, since sexual reproduction in the yeast experiment occurred only once every 30 generations. The effective number of independent observations underlying the empirical comparisons is therefore likely smaller than 1,161, and the null distributions and confidence intervals reported above should be interpreted as characterising sampling variability under an idealised independence assumption, rather than as fully accounting for the dependence structure among linked genomic windows.

## 5 Extracting multiallelic trajectories from pooled sequencing data

Multiallelic variation can be identified from next-generation sequencing (NGS) data using appropriate variant calling and filtering strategies. This includes the detection of small haplotypes (combinations of nearby SNPs) and structural variation, as well as estimation of allele frequencies in pooled sequencing samples. To illustrate this, we analyzed pooled sequencing data from a population of *Drosophila melanogaster* that was sampled at multiple time points. Variant calling was performed using FreeBayes (Garrison and Marth, 2012), which supports the identification of complex haplotypes and structural variants. Our analysis focused on chromosome arm 3R across three distinct sampled time points (July 2009, July 2012, and October 2015) from the “Linvilla” population, which is part of the “Drosophila in Time and Space” (DEST) project (Kapun et al., 2021). We applied a stringent filter, retaining only sites where each detected allele was supported by at least three reads at every sampled time point. This approach uncovered a wide range of multiallelic variation: we found 17,678 variant sites with three alleles, 1,602 with four, 180 with five, 14 with six, and 2 with seven alleles (see Figure S6). Considerable numbers of multiallelic variants were also found when we used different filtering thresholds (see Figure S7). We provide the code for this extraction pipeline in our github repository https://github.com/NikolasVellnow/selection_multiallelic.

**Figure S6.**
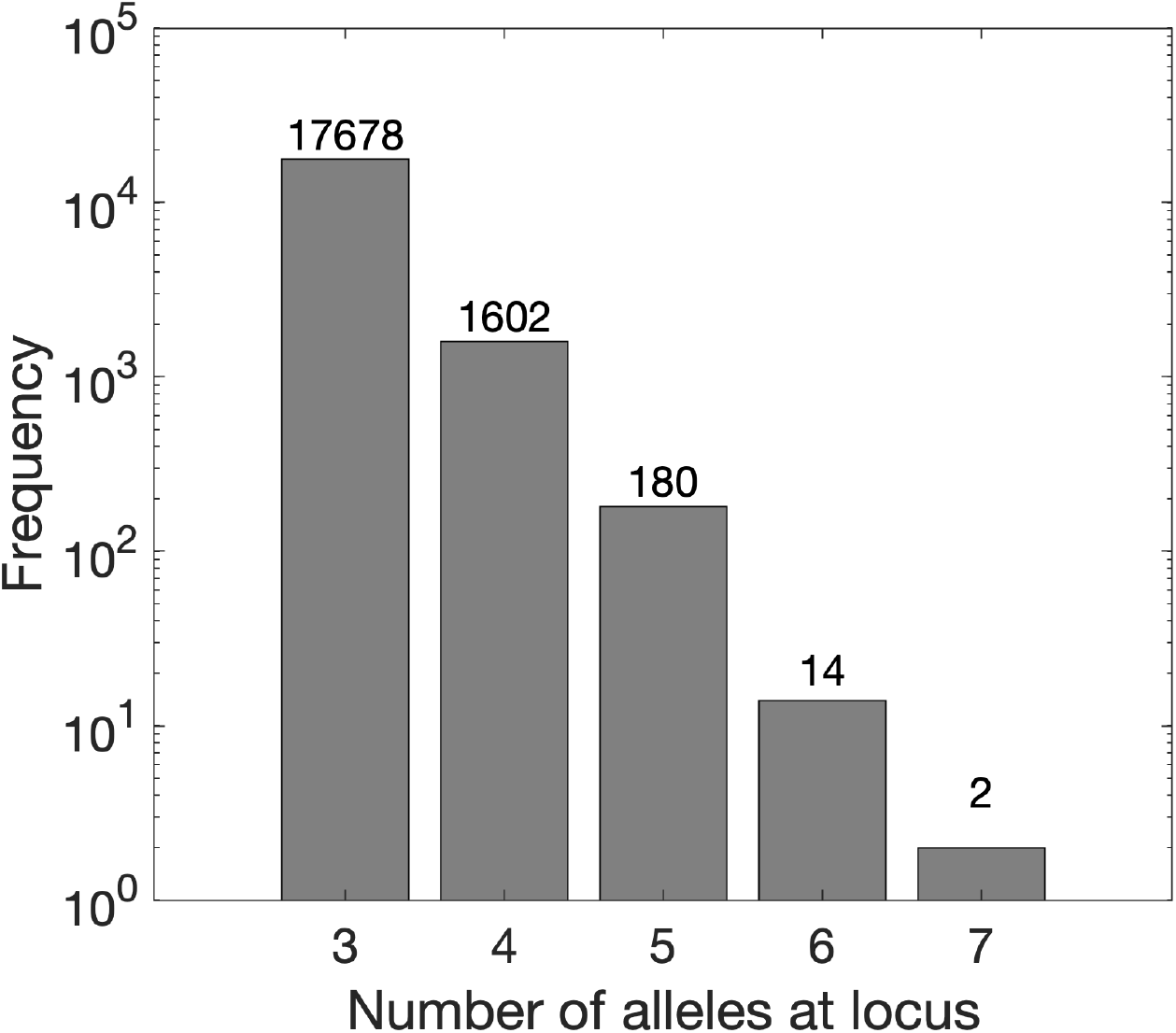
The number of sites that harbor more than two alleles in a natural D. melanogaster population. Data are shown for the chromosome arm 3R of the ‘Linvilla’ population obtained for three different time points (July 2009, July 2012, and October 2015), which is part of the DEST dataset (Kapun et al., 2021). We included only sites for which each allele was supported by at least three reads at each of the three sampled time points, i.e. we applied a hard filter.

Whether an allele is considered present depends on the filtering criteria applied to sequencing reads. In pooled sequencing data, a common approach is to require that each allele be supported by a minimum number of reads. In our primary analysis we required at least three reads supporting each allele at every sampled time point. Varying this threshold changes the number of detected multiallelic sites (see Figure S7). However, even under more stringent thresholds (e.g. requiring ≥ 7 reads per allele), substantial numbers of sites with three or four alleles remain detectable.

These results demonstrate that multiallelic polymorphisms are common in pooled sequencing data and can provide suitable trajectories for the inference framework developed in this study.

**Figure S7.**
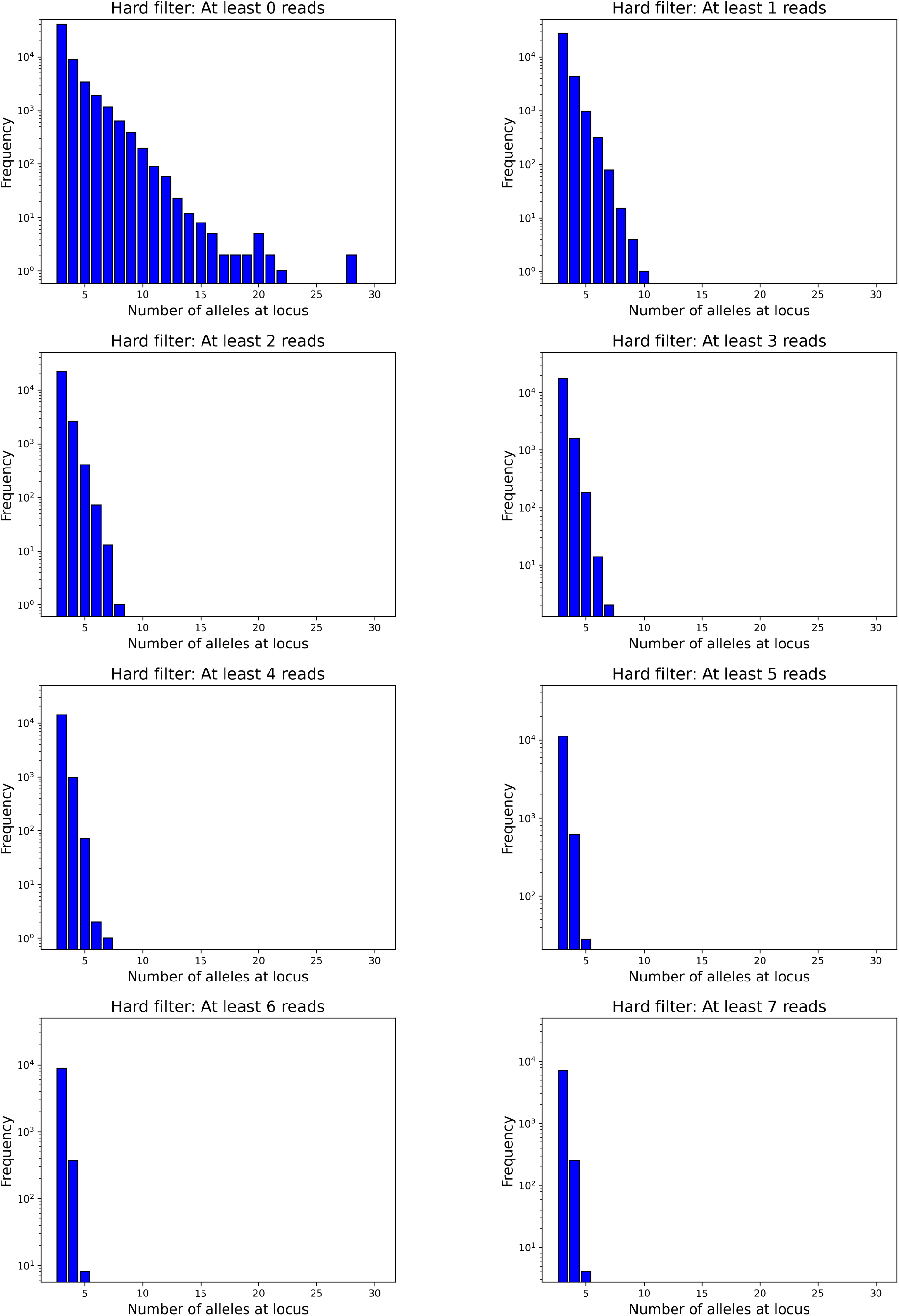
Sensitivity of number of alleles on hard filtering. Histograms are shown for the number of sites harbouring different numbers of alleles. The only sites included were those in which all alleles/haplotypes were supported by at least the specified minimum number of reads, at each of the three time points. This filtering criterion, for the minimum number of reads, was varied in the different subplots.

## 6 Probability of detection and thresholds

Here we provide calculations and simulations to substantiate some of the arguments given in the Discussion of the main text, concerning detection and threshold effects.

We used MATLAB for this purpose.

First, we modelled the probability of detection, for a coverage of *x*, as 1 minus the probability that the allele will not be contained in any of the *x* sequenced reads, i.e., probability of detection = 1 − (1 − *p*)^*x*^, where *p* is the relative frequency of the allele. We then plotted the probability of detection against the coverage for several values of *p* (see Figure S8a). Second, we simulated evolution of a locus with ten alleles, with a randomly generated **F** matrix. We then imitated the effect that a detection threshold would have on the allele frequency trajectories to be analyzed (see Figure S8b).

**Figure S8.**
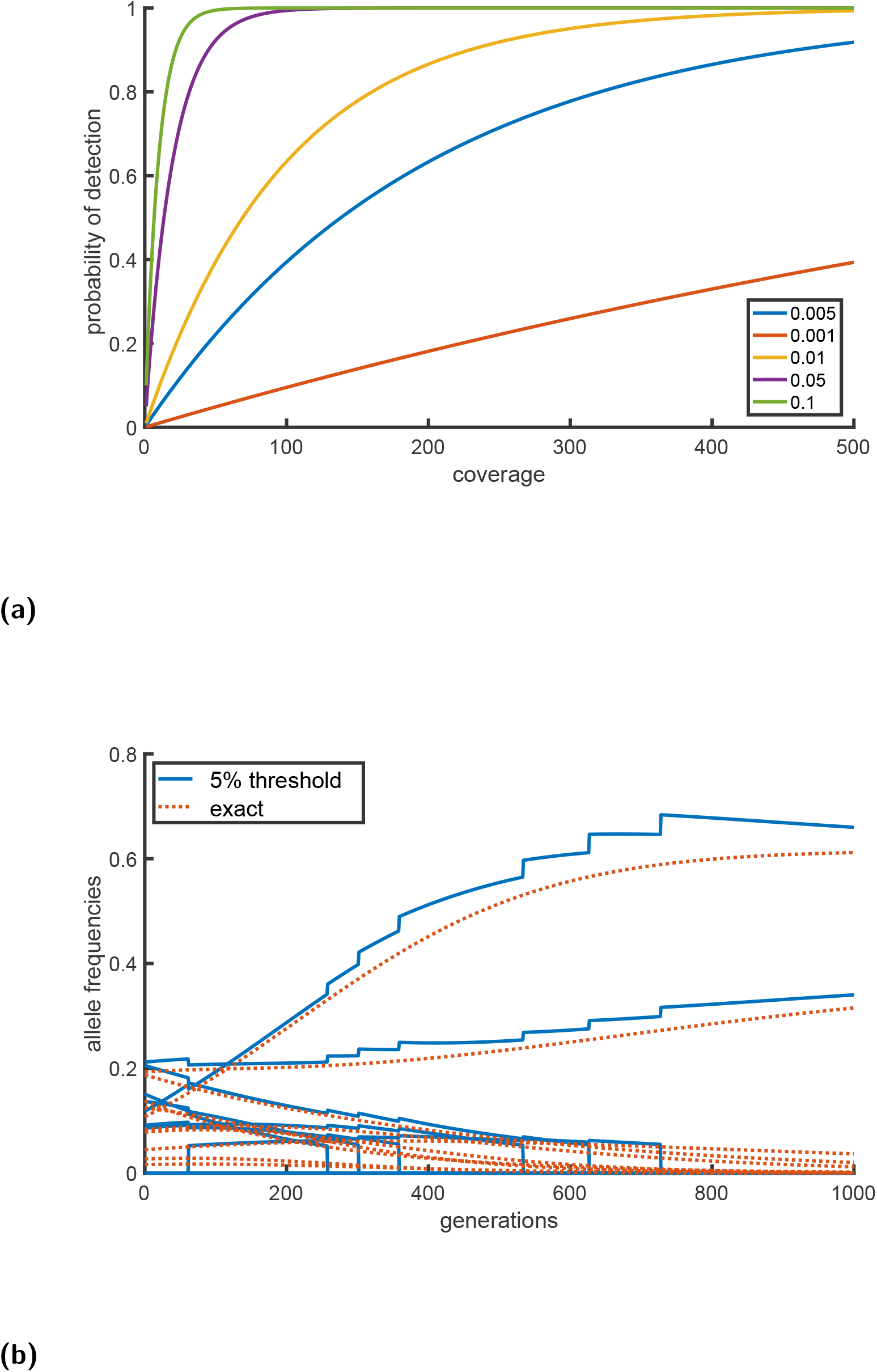
Illustration of observation thresholds in the analysis of multiallelic trajectories. (a) Probability to detect a genetic variant at a given frequency assuming different levels of sequencing coverage, as applied to pooled samples. (b) Plots of the exact allele frequency trajectories (red), and trajectories after applying a 5% threshold (blue) for a locus with ten alleles and an arbitrarily chosen scheme of selection. The threshold: (i) sets allele frequencies below 5% to zero, and (ii) normalizes the remaining frequencies so that they sum to unity. A consequence of (i) and (ii) are the ‘steps’ in the blue curves.

## Footnotes

1 We work within census population size, *N* , and not the effective population size, *N*_*e*_. However, incorporation of *N*_*e*_ into the Wright-Fisher model is possible (Zhao, Gossmann, and Waxman, 2016), and is common under the diffusion approximation.

2 With *µ*_*i,j*_ the probability of mutation from allele *A*_*j*_ to allele *A*_*i*_ (with *µ*_*i,i*_ = 0), the matrix **M** has the following form. For the elements *M*_*i,j*_ with *i* ≠ *j* (off diagonal elements), we have *M*_*i,j*_ = *µ*_*i,j*_ . In contrast, the diagonal elements are negative and given by 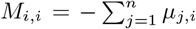. A consequence of this form of **M** is that all of its columns sum to zero.

3 Equation (6) is an approximation, based on the assumption that the effects of selection are small, so that on the right hand side, **D**(**x**) has been neglected in comparison with **x** (Ewens, 2004). When selection is not weak, **D**(**x**) cannot be neglected, for example, as occurs when there are lethal genotypes (Waxman and Overall, 2020).

4 An alternative and distinct derivation to the one we present in Appendix C, of the appearance of **V**(**x**) in selection, is via probabilistic reasoning (Barton and Turelli, 1987).

5 In the numerical results we present later we do not assume selection is weak, and we use the full form of **D**(**x**) given in Eq. (7).

6 In explicit calculations based on the diffusion approximation, the census size, *N* , is usually replaced by the effective population size, *N*_*e*_, to take into account deviations from an ideal Wright-Fisher model (Wright, 1931).

7 The model adopted for fluctuating selection, where *F*_*i,j*_ is given by Eq. (17), leads to a simple result for the averaged force of selection in terms the parameter *λ*, the matrix **V**(**x**), and the vector **x**. However, the model has the feature that the variance of fitness-effects of homozygotes is *twice* that of heterozygotes (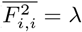, but 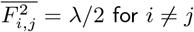)

8 A more general form of negative frequency-dependent selection is given by *F*_*i,j*_ (**x**) = −*c*_*i,j*_*x*_*i*_*x*_*j*_ with *c*_*i,j*_ = *c*_*j,i*_ and 0 ≤ *c*_*i,j*_ *<* 1.

9 We note that Eq. (27) is an approximation when fitnesses are not multiplicative, however for |*F*_*i,j*_ | ≪ 1 it is a good approximation (Nagylaki, 1992) (P. 252).

10 In the main text we wrote the equation for allele frequency changes as **X**(*t* + 1) = **X**(*t*) + **D**(**X**(*t*)). In this appendix we find it helpful to be more explicit, and we additionally list, as an argument of **D**, the **F** matrix that it depends upon.

11 ^11^Note that in the first equation, **X**(*T*_1_) is used on the right hand side, while in all subsequent equations **Y**’s are used.

## References

Aggeli, Dimitra et al. (June 2022). “Overdominant and Partially Dominant Mutations Drive Clonal Adaptation in Diploid Saccharomyces Cerevisiae”. In: Genetics 221.2, iyac061. issn: 1943-2631. doi: 10.1093/genetics/iyac061.

Akin, Ethan (1990). “The Differential Geometry of Population Genetics and Evolutionary Games”. In: Mathematical and Statistical Developments of Evolutionary Theory. Ed. by Sabin Lessard. Dordrecht: Springer Netherlands, pp. 1–93. isbn: 978-94-009-0513-9. doi: 10.1007/978-94-009-0513-9_1.

Axelsson, Erik et al. (Mar. 2013). “The Genomic Signature of Dog Domestication Reveals Adaptation to a Starch-Rich Diet”. In: Nature 495.7441, pp. 360–364. issn: 1476-4687. doi: 10.1038/nature11837.

Ayala, Francisco J. and Cathryn A. Campbell (1974). “Frequency-Dependent Selection”. In: Annual Review of Ecology and Systematics 5, pp. 115–138. issn: 0066-4162. JSTOR: 2096883.

Barton, N. H. and Michael Turelli (Apr. 1987). “Adaptive Landscapes, Genetic Distance and the Evolution of Quantitative Characters”. In: Genetics Research 49.2, pp. 157–173. issn: 1469-5073, 0016-6723. doi: 10.1017/S0016672300026951.

Burger, J. et al. (Mar. 2007). “Absence of the Lactase-Persistence-Associated Allele in Early Neolithic Europeans”. In: Proceedings of the National Academy of Sciences 104.10, pp. 3736–3741. doi: 10.1073/pnas.0607187104.

Bürger, Reinhard (Sept. 2000). The Mathematical Theory of Selection, Recombination, and Mutation. John Wiley & Sons. isbn: 978-0-471-98653-9.

Burke, Molly K., Gianni Liti, and Anthony D. Long (Jan. 2014). “Standing Genetic Variation Drives Repeatable Experimental Evolution in Outcrossing Populations of Saccharomyces Cerevisiae”. In: Molecular Biology and Evolution 31.12, pp. 3228–3239. issn: 0737-4038, 1537-1719. doi: 10.1093/molbev/msu256.

Caldon, Matteo et al. (2024). “Gelada Genomes Highlight Events of Gene Flow, Hybridisation and Local Adaptation That Track Past Climatic Changes”. In: Molecular Ecology n/a.n/a, e17514. issn: 1365-294X. doi: 10.1111/mec.17514.

Campbell, Ian M. et al. (Mar. 2016). “Multiallelic Positions in the Human Genome: Challenges for Genetic Analyses”. In: Human mutation 37.3, pp. 231–234. issn: 1059-7794. doi: 10.1002/humu.22944.

Charlesworth, B., M. T. Morgan, and D. Charlesworth (Aug. 1993). “The Effect of Deleterious Mutations on Neutral Molecular Variation”. In: Genetics 134.4, pp. 1289–1303. issn: 0016-6731. doi: 10.1093/genetics/134.4.1289.

Charlesworth, Deborah, Xavier Vekemans, et al. (2005). “Plant Self-Incompatibility Systems: A Molecular Evolutionary Perspective”. In: New Phytologist 168.1, pp. 61–69. issn: 1469-8137. doi: 10.1111/j.1469-8137.2005.01443.x.

Christiansen, Freddy Bugge (1999). Population Genetics of Multiple Loci. Chichester, New York, Weinheim, Brisbane, Singapore, Toronto: John Wiley & Sons Ltd. isbn: 978-0-471-97979-1.

Clark, Andrew G. (2004). “The Role of Haplotypes in Candidate Gene Studies”. In: Genetic Epidemiology 27.4, pp. 321–333. issn: 1098-2272. doi: 10.1002/gepi.20025.

Colbran, Laura L, Fabian C Ramos-Almodovar, and Iain Mathieson (June 2023). “A Gene-Level Test for Directional Selection on Gene Expression”. In: Genetics 224.2, iyad060. issn: 1943-2631. doi: 10.1093/genetics/iyad060.

Cornetti, Luca et al. (June 2024). “Long-Term Balancing Selection for Pathogen Resistance Maintains Trans-Species Polymorphisms in a Planktonic Crustacean”. In: Nature Communications 15.1, p. 5333. issn: 2041-1723. doi: 10.1038/s41467-024-49726-8.

Decaestecker, Ellen et al. (Dec. 2007). “Host–Parasite ‘Red Queen’ Dynamics Archived in Pond Sediment”. In: Nature 450.7171, pp. 870–873. issn: 1476-4687. doi: 10.1038/nature06291.

Dou, Yanmei et al. (July 2018). “Detecting Somatic Mutations in Normal Cells”. In: Trends in Genetics 34.7, pp. 545–557. issn: 0168-9525. doi: 10.1016/j.tig.2018.04.003.

Ebert, Dieter and Peter D. Fields (Dec. 2020). “Host–Parasite Co-Evolution and Its Genomic Signature”. In: Nature Reviews Genetics 21.12, pp. 754–768. issn: 1471-0064. doi: 10.1038/s41576-020-0269-1.

Edi, Constant V. et al. (Mar. 2014). “CYP6 P450 Enzymes and ACE-1 Duplication Produce Extreme and Multiple Insecticide Resistance in the Malaria Mosquito Anopheles Gambiae”. In: PLOS Genetics 10.3, e1004236. issn: 1553-7404. doi: 10.1371/journal.pgen.1004236.

Evans, Merran, N. A. J. Hastings, and Brian Peacock (June 1993). Statistical Distributions. Subsequent Edition. New York, NY: John Wiley & Sons Inc. isbn: 978-0-471-55951-1.

Ewens, Warren J. (Jan. 2004). Mathematical Population Genetics 1: Theoretical Introduction. 2nd ed. 2004 Edition. New York: Springer. isbn: 978-0-387-20191-7.

Fariello, María Inés et al. (Mar. 2013). “Detecting Signatures of Selection Through Haplotype Differentiation Among Hierarchically Structured Populations”. In: Genetics 193.3, pp. 929–941. issn: 1943-2631. doi: 10.1534/genetics.112.147231.

Faust, Karoline et al. (June 2015). “Metagenomics Meets Time Series Analysis: Unraveling Microbial Community Dynamics”. In: Current Opinion in Microbiology. Environmental Microbiology • Extremophiles 25, pp. 56–66. issn: 1369-5274. doi: 10.1016/j.mib.2015.04.004.

Fisher, RA (1922). “On the Dominance Ratio”. In: Proc. R. Soc. Edinburgh. Vol. 42, pp. 321–4.

Franssen, Susanne U., Nicholas H. Barton, and Christian Schlötterer (Jan. 2017). “Reconstruction of Haplotype-Blocks Selected during Experimental Evolution”. In: Molecular Biology and Evolution 34.1, pp. 174–184. issn: 0737-4038. doi: 10.1093/molbev/msw210.

Garcia Castillo Diego et al. (Oct. 2024). “Predicting Rapid Adaptation in Time from Adaptation in Space: A 30-Year Field Experiment in Marine Snails”. In: Science Advances 10.41, eadp2102. doi: 10.1126/sciadv.adp2102.

Garrison, Erik and Gabor Marth (July 2012). Haplotype-Based Variant Detection from Short-Read Sequencing. doi: 10.48550/arXiv.1207.3907. arXiv: 1207.3907.

Gillespie, John H. H. (Aug. 2004). Population Genetics: A Concise Guide. 2nd ed. Baltimore, Md.: Johns Hopkins University Press. isbn: 978-0-8018-8009-4.

Gossmann, Toni I, Megan Woolfit, and Adam Eyre-Walker (Dec. 2011). “Quantifying the Variation in the Effective Population Size Within a Genome”. In: Genetics 189.4, pp. 1389–1402. issn: 1943-2631. doi: 10.1534/genetics.111.132654.

Gossmann, Toni I., Peter D. Keightley, and Adam Eyre-Walker (Jan. 2012). “The Effect of Variation in the Effective Population Size on the Rate of Adaptive Molecular Evolution in Eukaryotes”. In: Genome Biology and Evolution 4.5, pp. 658–667. doi: 10.1093/gbe/evs027.

Gossmann, Toni I. and David Waxman (Apr. 2022). “Correcting Bias in Allele Frequency Estimates Due to an Observation Threshold: A Markov Chain Analysis”. In: Genome Biology and Evolution 14.4, evac047. issn: 1759-6653. doi: 10.1093/gbe/evac047.

Gossmann, Toni I., David Waxman, and Adam Eyre-Walker (Jan. 2014). “Fluctuating Selection Models and Mcdonald-Kreitman Type Analyses”. In: PLOS ONE 9.1, e84540. issn: 1932-6203. doi: 10.1371/journal.pone.0084540.

Haasl, Ryan J., Ross C. Johnson, and Bret A. Payseur (June 2014). “The Effects of Microsatellite Selection on Linked Sequence Diversity”. In: Genome Biology and Evolution 6.7, p. 1843. doi: 10.1093/gbe/evu134.

Haasl, Ryan J. and Bret A. Payseur (Feb. 2013). “Microsatellites as Targets of Natural Selection”. In: Molecular Biology and Evolution 30.2, pp. 285–298. issn: 1537-1719. doi: 10.1093/molbev/mss247.

Hartl, Daniel L. and Andrew G. Clark (1997). Principles of Population Genetics. Third edition. Hartl, Daniel L.; Harvard Univ., Cambridge, MA, USA: Sinauer Associates, Inc. isbn: 0-87893-306-9.

Hedrick, Philip W. (Dec. 2012). “What Is the Evidence for Heterozygote Advantage Selection?” In: Trends in Ecology & Evolution 27.12, pp. 698–704. issn: 0169-5347. doi: 10.1016/j.tree.2012.08.012.

Heyne, H. O. et al. (Jan. 2023). “Mono- and Biallelic Variant Effects on Disease at Biobank Scale”. In: Nature 613.7944, p. 519. doi: 10.1038/s41586-022-05420-7.

Higham, Nicholas J. (Jan. 2008). Functions of Matrices: Theory and Computation. Philadelphia, PA, USA: Society for Industrial and Applied Mathematics. isbn: 978-0-89871-646-7978-0-89871-777-8. doi: 10.1137/1.9780898717778.

Hildebrand, Falk, Toni I. Gossmann, et al. (July 2021). “Dispersal Strategies Shape Persistence and Evolution of Human Gut Bacteria”. In: Cell Host & Microbe 29.7, 1167–1176.e9. issn: 1931-3128. doi: 10.1016/j.chom.2021.05.008.

Hildebrand, Falk, Lucas Moitinho-Silva, et al. (Oct. 2019). “Antibiotics-Induced Monodominance of a Novel Gut Bacterial Order”. In: Gut 68.10, pp. 1781–1790. issn: 0017-5749, 1468-3288. doi: 10.1136/gutjnl-2018-317715.

Hoerl, Arthur E. and Robert W. Kennard (Feb. 1970). “Ridge Regression: Biased Estimation for Nonorthogonal Problems”. In: Technometrics 12.1, pp. 55–67. issn: 0040-1706. doi: 10.1080/00401706.1970.10488634.

Huerta-Sanchez, Emilia, Rick Durrett, and Carlos D Bustamante (Jan. 2008). “Population Genetics of Polymorphism and Divergence Under Fluctuating Selection”. In: Genetics 178.1, pp. 325–337. issn: 1943-2631. doi: 10.1534/genetics.107.073361.

Jensen, Louis and Edward Pollak (1969). “Random Selective Advantages of a Gene in a Finite Population”. In: Journal of Applied Probability 6.1, pp. 19–37. issn: 0021-9002. doi: 10.2307/3212274. JSTOR: 3212274.

Kapun, Martin et al. (Dec. 2021). “Drosophila Evolution over Space and Time (DEST): A New Population Genomics Resource”. In: Molecular Biology and Evolution 38.12, pp. 5782–5805. issn: 1537-1719. doi: 10.1093/molbev/msab259.

Kimura, Motoo (1955). “Random Genetic Drift in Multi-Allelic Locus”. In: Evolution 9.4, pp. 419–435. issn: 0014-3820. doi: 10.2307/2405476.JSTOR: 2405476.

Kimura, Motoo and James F. Crow (Apr. 1964). “The Number of Alleles That Can Be Maintained in a Finite Population”. In: Genetics 49.4, pp. 725–738. issn: 0016-6731.

Korneliussen, Thorfinn Sand, Anders Albrechtsen, and Rasmus Nielsen (Nov. 2014). “ANGSD: Analysis of Next Generation Sequencing Data”. In: BMC Bioinformatics 15.1, p. 356. issn: 1471-2105. doi: 10.1186/s12859-014-0356-4.

Kosel, Brinja et al. (2024). “Evolved Readers of 5-Carboxylcytosine CpG Dyads Reveal a High Versatility of the Methyl-CpG-Binding Domain for Recognition of Noncanonical Epigenetic Marks”. In: Angewandte Chemie International Edition 63.17, e202318837. issn: 1521-3773. doi: 10.1002/anie.202318837.

Latter, B. D. H. and C. E. Novitski (Aug. 1969). “Selection in Finite Populations with Multiple Alleles I. Limits to Directional Selection”. In: Genetics 62.4, p. 859. doi: 10.1093/genetics/62.4.859.

Lewin, Loveday and Adam Eyre-Walker (2026). “A Comparative Analysis of Long-Term Effective Population Sizes Across Eukaryotes”. In: Molecular Ecology 35.4, e70265. issn: 1365-294X. doi: 10.1111/mec.70265.

Lewontin, R C,R R Ginzburg, and S D Tuljapurkar (Jan. 1978). “HETEROSIS AS AN EXPLANATION FOR LARGE AMOUNTS OF GENIC POLYMORPHISM”. In: Genetics 88.1, pp. 149–169. issn: 1943-2631. doi: 10.1093/genetics/88.1.149.

Liu, Jie and Xuehua Zhong (Feb. 2024). “Epiallelic Variation of Non-Coding RNA Genes and Their Phenotypic Consequences”. In: Nature Communications 15.1, p. 1375. issn: 2041-1723. doi: 10.1038/s41467-024-45771-5.

Lupski, James R. (July 2013). “Genome Mosaicism—One Human, Multiple Genomes”. In: Science 341.6144, pp. 358–359. doi: 10.1126/science.1239503.

Malaspinas, Anna-Sapfo (2016). “Methods to Characterize Selective Sweeps Using Time Serial Samples: An Ancient DNA Perspective”. In: Molecular Ecology 25.1, pp. 24–41. issn: 1365-294X. doi: 10.1111/mec.13492.

McKenna, Aaron et al. (Jan. 2010). “The Genome Analysis Toolkit: A MapReduce Framework for Analyzing next-Generation DNA Sequencing Data”. In: Genome Research 20.9, pp. 1297–1303. issn: 1088-9051, 1549-5469. doi: 10.1101/gr.107524.110.

Nagylaki, Thomas (June 1992). Introduction to Theoretical Population Genetics. 1st ed. Berlin Heidelberg: Springer. isbn: 978-3-540-53344-3.

Nesta, Alex V., Denisse Tafur, and Christine R. Beck (Aug. 2021). “Hotspots of Human Mutation”. In: Trends in Genetics 37.8, pp. 717–729. issn: 0168-9525. doi: 10.1016/j.tig.2020.10.003.

Nielsen, Rasmus, Andrew H. Vaughn, and Yun Deng (Jan. 2025). “Inference and Applications of Ancestral Recombination Graphs”. In: Nature Reviews Genetics 26.1, pp. 47–58. issn: 1471-0064. doi: 10.1038/s41576-024-00772-4.

Ollivier, Morgane et al. (Nov. 2016). “Amy2B Copy Number Variation Reveals Starch Diet Adaptations in Ancient European Dogs”. In: Royal Society Open Science 3.11, p. 160449. doi: 10.1098/rsos.160449.

Oota, Satoshi (Apr. 2020). “Somatic Mutations – Evolution within the Individual”. In: Methods. RNA-Seq: Methods and Applications 176, pp. 91–98. issn: 1046-2023. doi: 10.1016/j.ymeth.2019.11.002.

Overall, A. D. J. and D. Waxman (Aug. 2024). “Influence of Selection on the Probability of Fixation at a Locus with Multiple Alleles”. In: BMC Genomics 25.1, p. 819. issn: 1471-2164. doi: 10.1186/s12864-024-10733-0.

Paudel, Yogesh et al. (July 2013). “Evolutionary Dynamics of Copy Number Variation in Pig Genomes in the Context of Adaptation and Domestication”. In: BMC Genomics 14.1, p. 449. issn: 1471-2164. doi: 10.1186/1471-2164-14-449.

Perry, George H. et al. (Oct. 2007). “Diet and the Evolution of Human Amylase Gene Copy Number Variation”. In: Nature Genetics 39.10, pp. 1256–1260. issn: 1546-1718. doi: 10.1038/ng2123.

Peters, Jo (Aug. 2014). “The Role of Genomic Imprinting in Biology and Disease: An Expanding View”. In: Nature Reviews Genetics 15.8, pp. 517–530. issn: 1471-0064. doi: 10.1038/nrg3766.

Phillips, Mark A. et al. (2020). “Increased Time Sampling in an Evolve-and-Resequence Experiment with Outcrossing Saccharomyces Cerevisiae Reveals Multiple Paths of Adaptive Change”. In: Molecular Ecology 29.24, pp. 4898–4912. issn: 1365-294X. doi: 10.1111/mec.15687.

Radwan, Jacek et al. (Apr. 2020). “Advances in the Evolutionary Understanding of MHC Polymorphism”. In: Trends in Genetics 36.4, pp. 298–311. issn: 0168-9525. doi: 10.1016/j.tig.2020.01.008.

Rhoads, Anthony and Kin Fai Au (Oct. 2015). “PacBio Sequencing and Its Applications”. In: Genomics, Proteomics & Bioinformatics 13.5, pp. 278–289. issn: 1672-0229. doi: 10.1016/j.gpb.2015.08.002.

Ruzicka, Filip et al. (2025). “A Century of Theories of Balancing Selection”. In: Biological Reviews. issn: 1469-185X. doi:10.1111/brv.70103.

Schlötterer, C. et al. (May 2015). “Combining Experimental Evolution with Next-Generation Sequencing: A Powerful Tool to Study Adaptation from Standing Genetic Variation”. In: Heredity 114.5, pp. 431–440. issn: 1365-2540. doi: 10.1038/hdy.2014.86.

Sedlazeck, Fritz J. et al. (June 2018). “Piercing the Dark Matter: Bioinformatics of Long-Range Sequencing and Mapping”. In: Nature Reviews Genetics 19.6, pp. 329–346. issn: 1471-0064. doi: 10.1038/s41576-018-0003-4.

Sellis, Diamantis et al. (July 2016). “Heterozygote Advantage Is a Common Outcome of Adaptation in Saccharomyces Cerevisiae”. In: Genetics 203.3, pp. 1401–1413. issn: 1943-2631. doi: 10.1534/genetics.115.185165.

Shahshahani, S. (1979). A New Mathematical Framework for the Study of Linkage and Selection. American Mathematical Soc. isbn: 978-0-8218-2211-1.

Sharma, N. and G. R. Cutting (Mar. 2020). “The Genetics and Genomics of Cystic Fibrosis”. In: Journal of Cystic Fibrosis. ECFS Cystic Fibrosis Research 19, S5–S9. issn: 1569-1993. doi: 10.1016/j.jcf.2019.11.003.

Siljestam, Mattias and Claus Rueffler (Nov. 2024). “Heterozygote Advantage Can Explain the Extraordinary Diversity of Immune Genes”. In: eLife 13. Ed. by Yaroslav Ispolatov, Detlef Weigel, and Yaroslav Ispolatov, e94587. issn: 2050-084X. doi: 10.7554/eLife.94587.

Smith, J. M. and J. Haigh (Feb. 1974). “The Hitch-Hiking Effect of a Favourable Gene”. In: Genetical Research 23.1, pp. 23–35.

Spencer, H. G. and R. W. Marks (Oct. 1988). “The Maintenance of Single-Locus Polymorphism. I. Numerical Studies of a Viability Selection Model”. In: Genetics 120.2, pp. 605–613. issn: 0016-6731. doi: 10.1093/genetics/120.2.605.

Tajima, F. (Nov. 1989). “Statistical Method for Testing the Neutral Mutation Hypothesis by DNA Polymorphism”. In: Genetics 123.3, pp. 585–595. issn: 0016-6731.

Takayama, Seiji and Akira Isogai (June 2005). “SELF-INCOMPATIBILITY IN PLANTS”. In: Annual Review of Plant Biology 56.Volume 56, 2005, pp. 467–489. issn: 1543-5008, 1545-2123. doi: 10.1146/annurev.arplant.56.032604.144249.

The Mathworks Inc. (2025). MATLAB. The MathWorks Inc. Natick, Massachusetts, United States.

Tuckwell, Henry C. (Dec. 2019).Elementary Applications of Probability Theory: With an Introduction to Stochastic Differential Equations. 2nd edition. Chapman and Hall/CRC. isbn: 978-0-367-44905-6.

Vellnow, Nikolas, Toni I. Gossmann, and David Waxman (Apr. 2024). “The Pseudoentropy of Allele Frequency Trajectories, the Persistence of Variation, and the Effective Population Size”. In: BioSystems 238, p. 105176. issn: 0303-2647. doi: 10.1016/j.biosystems.2024.105176.

Wang, Yiguan and Darren J Obbard (Aug. 2023). “Experimental Estimates of Germline Mutation Rate in Eukaryotes: A Phylogenetic Meta-Analysis”. In: Evolution Letters 7.4, pp. 216–226. issn: 2056-3744. doi: 10.1093/evlett/qrad027.

Wang, Yunhao, Yue Zhao, et al. (Nov. 2021). “Nanopore Sequencing Technology, Bioinformatics and Applications”. In: Nature Biotechnology 39.11, pp. 1348–1365. issn: 1546-1696. doi: 10.1038/s41587-021-01108-x.

Watterson, G. A. (Apr. 1975). “On the Number of Segregating Sites in Genetical Models without Recombination”. In: Theoretical Population Biology 7.2, pp. 256–276. issn: 0040-5809. doi: 10.1016/0040-5809(75)90020-9.

Waxman, David and Andrew D. J. Overall (Apr. 2020). “Influence of Dominance and Drift on Lethal Mutations in Human Populations”. In: Frontiers in Genetics 11. issn: 1664-8021. doi: 10.3389/fgene.2020.00267.

Whitehouse, Logan S and Daniel R Schrider (July 2023). “Timesweeper: Accurately Identifying Selective Sweeps Using Population Genomic Time Series”. In: Genetics 224.3, iyad084. issn: 1943-2631. doi: 10.1093/genetics/iyad084.

Williams, George Christopher (1966). Adaptation and Natural Selection: A Critique of Some Current Evolutionary Thought. Princeton University Press. isbn: 978-0-691-18550-7.

Wortel, Meike T. et al. (2023). “Towards Evolutionary Predictions: Current Promises and Challenges”. In: Evolutionary Applications 16.1, pp. 3–21. issn: 1752-4571. doi: 10.1111/eva.13513.

Wright, Sewall (Mar. 1931). “Evolution in Mendelian Populations”. In: Genetics 16.2, pp. 97–159. issn: 0016-6731. doi: 10.1093/genetics/16.2.97.

Wright, Sewall (June 1939). “The Distribution of Self-Sterility Alleles in Populations”. In: Genetics 24.4, p. 538. doi: 10.1093/genetics/24.4.538.

Yamamoto, Fumiichiro (Sept. 2021). “Molecular Genetics and Genomics of the ABO Blood Group System”. In: Annals of Blood 6.0. issn: 2521-361X. doi: 10.21037/aob-20-71.

Yokoyama, S. and M. Nei (Mar. 1979). “Population Dynamics of Sex-Determining Alleles in Honey Bees and Self-Incompatibility Alleles in Plants”. In: Genetics 91.3, pp. 609–626. issn: 0016-6731. doi: 10.1093/genetics/91.3.609.

Zhao, Lei, Toni I. Gossmann, and David Waxman (Mar. 2016). “A Modified Wright–Fisher Model That Incorporates Ne: A Variant of the Standard Model with Increased Biological Realism and Reduced Computational Complexity”. In: Journal of Theoretical Biology 393, pp. 218–228. issn: 0022-5193. doi: 10.1016/j.jtbi.2016.01.002.

